# Multiple-testing corrections in case-control studies using identity-by-descent segments

**DOI:** 10.1101/2025.07.03.663057

**Authors:** Seth D. Temple, Nicola H. Chapman, Seung Hoan Choi, Anita L. DeStefano, Timothy A. Thornton, Ellen M. Wijsman, Elizabeth E. Blue

**Affiliations:** Department of Statistics, University of Washington, Seattle, Washington, USA; Department of Statistics, University of Michigan, Ann Arbor, Michigan, USA; Michigan Institute for Data Science, University of Michigan, Ann Arbor, Michigan, USA; Department of Biostatistics, Boston University School of Public Health, Boston, Massachusetts, USA; Division of Medical Genetics, Department of Medicine, University of Washington, Seattle, Washington, USA; Regeneron Genetics Center, Tarrytown, New York, USA; Department of Biostatistics, University of Washington, Seattle, Washington, USA; Department of Genome Sciences, University of Washington, Seattle, Washington, USA; Institute for Public Health Genetics, University of Washington, Seattle, Washington, USA; Brotman Bay Institute, Seattle, Washington, USA

**Keywords:** multiple testing, binary traits, Alzheimer’s disease, identity by descent, haplotypes, mean-reverting processes

## Abstract

Identity-by-descent (IBD) mapping provides complementary signals to genome-wide association studies (GWAS) when multiple causal haplotypes or variants are present, but not directly tested. However, failing to correct for multiple testing in case-control studies using IBD segments can lead to false discoveries. We propose the difference between case-case and control-control IBD rates as an IBD mapping statistic. For our hypothesis test, we use a computationally efficient approach from the stochastic processes literature to derive genome-wide significance levels that control the family-wise error rate (FWER). Whole genome simulations indicate that our method conservatively controls the FWER. Because positive selection can lead to false discoveries, we pair our IBD mapping approach with a selection scan so that one can contrast results for evidence of confounding due to recent sweeps or other mechanisms, like population structure, that increase IBD sharing. We developed automated and reproducible workflows to phase haplotypes, call local ancestry probabilities, and perform the IBD mapping scan, the former two tasks being important preprocessing steps for haplotype analyses. We applied our methods to search for Alzheimer’s disease (AD) risk loci in the Alzheimer’s Disease Sequencing Project (ADSP) genome data. We identified six genome-wide significant signals of AD risk among samples genetically similar to African and European reference populations and self-identified Amish samples. Variants in the six potential risk loci we detected have previously been associated with AD, dementia, and memory decline. Three genes at two potential risk loci have already been nominated as therapeutic targets. Overall, our scalable approach makes further use of large consortia resources, which are expensive to collect but provide insights into disease mechanisms.

**Highlights:**

- We propose a computationally efficient method to address multiple testing when scanning along the genome for differences in identity-by-descent rates of case-case and control-control pairs.
- Whole genome simulations indicate that our method conservatively controls the desired family-wise error rate.
- We performed three case-control scans from ancestry cohorts in the Alzheimer’s Disease Sequencing Project, detecting six genome-wide significant signals around potential risk loci.
- We show that positive selection can confound IBD mapping tests in samples genetically similar to Europeans.

## 1. Introduction

Population-based identity-by-descent (IBD) mapping complements a single variant GWAS, which is better suited for common variant associations [1, 2]. Genome-wide association studies (GWAS) and family-based studies have limited power to detect risk or causal variants that are recent and rare, particularly when those variants are observed in separate families [3]. IBD mapping encompasses methods that investigate the relationships between IBD haplotype segments and pheno-types. Such approaches can yield as much as a 40% power gain over GWAS when between 1 to 10% of haplotypes carry a causal variant [4].

Currently, there are few IBD mapping methods for binary and quantitative trait phenotypes. Albrechtsen et al. [5] examined the difference in genetic linkage within affected versus within unaffected individuals. Browning and Thompson [4] instead considered the ratio of the rates of long IBD segments detected between case-case pairs and case-control pairs. Gusev et al. [6] proposed an association test with IBD haplotype clusters, and Cai and Browning [7] and Chen et al. [8] proposed different variance component tests to find quantitative trait associations with IBD haplotypes. These IBD mapping methods have successfully detected associations between rare haplotypes and complex traits, including Parkinson’s disease [9], multiple sclerosis [10], amyotrophic lateral sclerosis [11], collagen diseases [12], serum triglycerides [13], systolic blood pressure [7], and type 1 diabetes [4].

Analyses of imputed variants, whole genome sequences (WGS), and inferred genealogies are alternative approaches to uncovering the etiologies of rare diseases. Many rare variant methods aggregate the evidence of multiple rare variants via burden tests [14, 15], weighted sums [16], or variance component tests [17]. Combining the signals of different rare variant tests into omnibus tests can also be powerful [18, 19]. Promising new methods use variance component tests to associate complex traits with unobserved ultra-rare variants placed on the branches of an inferred genealogical tree [20, 21, 22]. IBD mapping can outperform rare variant tests by testing for haplotype associations, where there may be multiple causal variants, some of which could be ungenotyped. It also does not involve defining a testing unit, such as gene boundaries [7].

Stolyarova et al. [23] argued that large-effect variants at rare to low frequencies are shared IBD from extended families, not ancestry groups, and so focusing on ancestry-specific variants [24, 25] may also be a less effective approach to disease mapping than IBD mapping. When the genetic disease comes from a low-frequency variant, Voight and Pritchard [26] explained how ascertaining individuals with the disease case phenotype induces cases that are more related to each other than controls are, and thereby excess cryptic relatedness. Voight and Pritchard [26] remarked that such ascertainment can lead to substantial confounding in single-variant GWAS under special conditions. Taking a different perspctive, the hypothesis test we propose in this paper leverages the signal of excess relatedness among case individuals to identify low frequency disease haplotypes.

It can be challenging in complex haplotype analyses to determine an appropriate genome-wide significance level. Since haplotype clusters can span considerable genetic distances, haplotype-based tests can be highly correlated. Using the GWAS significance level p ¡ 5e-8 in haplotype-based tests would seriously diminish power. For reference, Temple [27] and Temple and Browning [28] review several multiple-testing paradigms in statistical and population genetics. Here, we focus on the family-wise error rate (FWER), which is the probability of rejecting the null hypothesis one or more times when the null hypothesis is true [29].

The initial IBD mapping methods employed computationally intensive permutation tests to determine the FWER [30, 4, 6]. For instance, the Browning and Thompson [4] permutation test requires tens of GBs of memory and many hours to analyze a case-control cohort of a few thousand samples. By modeling transitions between IBD to non-IBD states as a Markov process, Browning and Thompson [4] proposed a genome-wide significance level that can be derived analytically. How-ever, their analytical approach involves numerous approximations [31, 32], assumes a constant population size, and requires prior knowledge about the coalescent times of the most recent common ancestors from whom the IBD alleles originated. In practice, studies using the Browning and Thompson [4] pairwise IBD rate ratio test choose the permutation test over the analytical approximation [10, 4].

When the correlated tests can be modeled as a parametric random process, we may be able to derive the genome-wide significance level from the properties of the stochastic process. Recently, Feingold et al. [33], Grinde et al. [34], Temple and Browning [28], and Cai and Browning [7] proposed model-based multiple-testing corrections for linkage analysis, admixture mapping, selection scans, and IBD mapping, respectively. These approaches are based on the Ornstein-Uhlenbeck (OU) process. They are, therefore, most valid when the test statistic is (asymptotically) normally distributed and the correlations of test statistics decline exponentially. The Temple and Browning [28] method detects regions with excess IBD rates. Following Temple and Browning [28], we will scan for regions where the difference in IBD rates between case-case and control-control pairs is extreme relative to the genome-wide central tendency. Compared to the Temple and Browning [28] scan for recent positive selection, sample ascertainment and classification of case/control status serve as the selection process.

For the pairwise IBD rate difference test, we propose analytical and simulation-based significance thresholds from an estimated OU process model. We show that the adjusted significance thresholds conservatively control the FWER under some central limit theorem conditions [35] and offer more statistical power than a naive Bonferroni correction. The pairwise IBD rate difference scan is automated and computationally efficient, allowing for the analysis of many binary phenotypes in biobanks representing hundreds of thousands of individuals. We applied our method to study Alzheimer’s disease (AD) in samples representing European, African, and Amish ancestry from the Alzheimer’s Disease Sequencing Project (ADSP).

## 2. Materials and Methods

### 2.1 Hypothesis testing framework

We define the difference in IBD rates test with the mathematical notation of Temple and Thompson [35] and Temple and Browning [28]. Let the case and control IBD rates overlapping the *m*^th^ focal position be 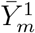 and 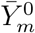. The hypothesis test we consider is

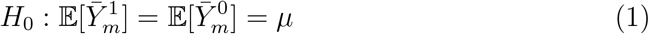

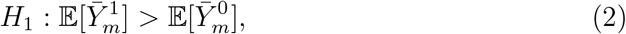

where *µ* is a genome-wide mean IBD rate around a locus. This null model is consistent with case and control sample sets having the same expected IBD rate. The alternative model is consistent with case haplotypes having a higher expected IBD rate, though the null model could also be rejected due to technical artifacts in IBD segment detection or other biological mechanisms.

The proposed hypothesis test (Equations 1 and 2) is similar to the test for selection described by Temple and Browning [28], where the null model is that the IBD rates 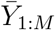 of a single sample set have the same genome-wide mean and that strong positive selection is an alternative model where some IBD rates are elevated. The two tests are intended to serve as examples of one-sided one and two-sample z tests.

For *M* IBD rates along the genome, let 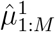 and 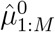 be the genome-wide average case and control IBD rates 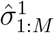 and 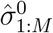 the genome-wide case and control standard deviations. The standardized case and control IBD rates are then 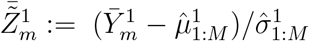 and 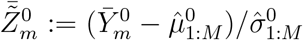. Under asymptotic conditions on sample sizes (more than a thousand), population sizes (more than ten thousand), and the detection threshold, the standardized IBD rates 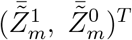 around the *m*^th^ locus converge weakly to the multivariate normally distribution [35], where *T* is the matrix transpose. Consequenly, the difference of the standardized IBD rates 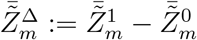 is asymptotically normally distributed [36].

We frame our test in terms of the standardized difference of IBD rates. Let 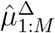 and 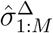 be the genome-wide average and standard deviation of the standardized differences. Then, the test is

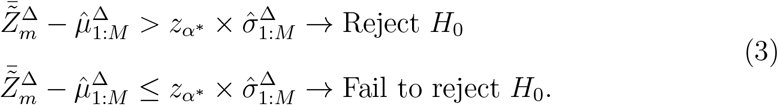

The significance level *α*^*^ comes from a multiple-testing correction at the family-wise significance level *α*, and 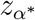 is the corresponding standard normal quantile. The IBD rate difference 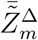 is distinct from the test statistic in Browning and Thompson [4], which is essentially the IBD rates ratio 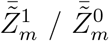.

### 2.2 An analytical approach to multiple testing

We desire multiple-testing corrections that adapt to the number of hypothesis tests we make. The Bonferroni correction would be very conservative because tests of markers in linkage disequilibrium (LD) are not independent. Similar to Temple and Browning [28], we model the standardized differences in standardized IBD rates along the genome as a correlated OU process. The multivariate OU process is normally distributed at every point, is spatially homogeneous, has the first-order Markov property, and each marginal, one-dimensional process is also an OU process. Temple and Browning [28] have shown that the OU assumptions of IBD rates are reasonable for some human genetics studies. We will treat the 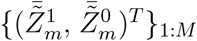 as a two-dimensional OU process and therefore 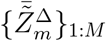 as a one-dimensional OU process.(Technically, 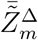 has an asymptotic variance of 2 [35, 37]; we will standardize the standardized differences to have a variance of 1 and, in good faith, not introduce further notation.)

Holding the genetic distance between consecutive focal positions to be constant *δ*, the covariance between standardized differences 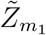 and 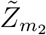 at different loci is

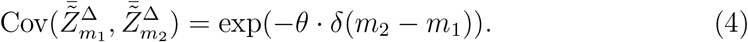

The exponential decay parameter *θ* captures the correlation between test statistics, which is unknown for the IBD rate difference process but can be estimated.

We employ the same analytical technique to control the FWER as Temple and Browning [28], which is an approximation method based on Siegmund and Yakir [38] and Feingold et al. [33]. Namely, for total genome length *L* (in Morgans), *C* chromosomes, and the Gaussian cumulative and density functions Φ and *ϕ*,

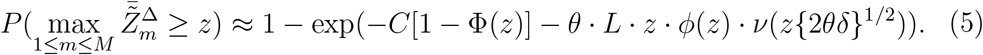

The function *ν*(·) accommodates the discretization of tests because the OU model is a continuous stochastic process. Hence, we will refer to this method as a *discrete-spacing analytical approach*. We use a root solver to determine 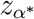 from Equation 5.

In Appendix A.2, we provide a *simulation-based approach* to control the FWER with our OU model. Under that approach, we treat 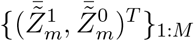 as OU processes with exponential decay parameters *θ*_1_ and *θ*_0_ and cross-correlation term *ρ*. We perform hundreds to thousands of simulations to determine the (1 − *α*)^th^ quantile of the simulated maxima of 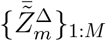. For most values of *θ*, the simulation-based and analytical approaches yield nearly identical thresholds. Still, the whole-genome simulation-based approach can take as long as ten minutes on an Intel 2.60 GHz core processing unit (CPU), compared to the instantaneous calculation in the analytical approach [27, 28].

### 2.3 Estimator of the exponential decay parameter

Before standardizing the case and control IBD rates, we adjust for extreme outliers that could be present in real genetic data; for example, positive selection can result in excess IBD rates, and low mappability or low marker density can result in deflated IBD rates [39, 40, 28]. First, we compute the initial case and control median IBD rates plus four times their standard deviations. Second, we compute revised case and control mean IBD rates and standard deviations, excluding the genetic positions where either case or control IBD rates exceed their initial median plus four standard deviations thresholds. We standardize the case and control IBD rates with the revised means and standard deviations. Third, we standardize the difference between the standardized case and control IBD rates.

We regress estimated autocovariances on genetic position to estimate the exponential decay parameter *θ* [28]. Given a recombination map, we use linear interpolation to hold the spacings between IBD rates constant. Then, we estimate the covariance between standardized IBD rate differences at genetic positions Δ times some integer constant apart, excluding positions where case or control IBD rates exceed their initial thresholds. We increment the integer scalars by one until the maximum difference between autocovariances is 4.0 centiMorgan (cM) apart. We fit a simple log-linear model with no intercept, where the integer-scaled Δ’s are the covariates and the estimated autocovariances are the response variables. The fitted slope parameter of the regression model 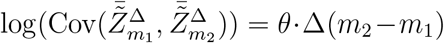 is an estimator 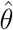 of the exponential decay parameter. Estimating *θ* takes less than a minute on an Intel 2.60 GHz CPU; most of the computing time is reading in the IBD segments.

### 2.4 Simulating IBD rate processes

#### 2.4.1 The case phenotype has no effect

To assess if our method controls the FWER, we used `msprime` [41] and `tskibd` [42] to simulate the ≥ 2.0 and ≥ 3.0 cM IBD rates of 2500 samples, randomly split into 1250 cases and 1250 controls. We explored different detection thresholds because, in the related study by Temple and Browning [28], the detection threshold impacts FWER and power. For each simulation, we created ten 100 cM chromosomes (10 Morgans in total) with a constant recombination rate of 1e-8. FWER is calculated as the percentage of the 500 null model simulations with at least one significant result. We considered human-like demographic scenarios, including a population bottleneck, three phases of exponential growth, and a constant population of 50,000 individuals [27, 35, 43, 40]. The demographic scenario affects the exponential decay parameter *θ* [28].

#### 2.4.2 Selective sweeps as a confounding model

Using the average genome-wide significance level of our simulations, we estimated the probability that we reject the null hypothesis in hard selective sweeps. This evolutionary scenario concerns an advantageous allele that increases in frequency as a function of a selection coefficient *s*. (See Temple [27] for a description of the alternative model.) IBD rates overlapping a selected locus were simulated with the Temple et al. [43] algorithm. We randomly assigned the 2500 diploid individuals into 1250 cases and 1250 controls, which inserts randomness into whether more or fewer cases than controls have the beneficial allele. Because cases and controls are expected to have the same counts of the selected allele, rejecting the null model means that positive selection has confounded our test.

### 2.5 Sequence data from the Alzheimer’s Disease Sequencing Project

AD is highly heritable and has a complex genetic architecture [44, 45]. There are rare variants underlying autosomal dominant AD (*APP, PSEN1*, and *PSEN2* ; OMIM: 104760, 104311, 600759) [46], rare variants with large effect sizes (e.g., *TREM2* ; OMIM: 605086), common variants with large and ancestry-specific effect sizes in the apolipoprotein E gene (*APOE*, OMIM: 107741) [44, 47, 48], and dozens of common variants with modest effect sizes [49, 50]. Most AD GWAS, including those by the ADSP, have focused on single nucleotide polymorphism (SNP) tests or SNP-by-SNP interactions [51, 52]. We applied our IBD mapping method to find rare to low frequency shared haplotypes enriched in AD cases relative to AD controls.

We used the autosomes of the ADSP whole genome sequence data (release 4) [53], with variant calling and quality controls performed from the Leung et al. [54] pipeline. We analyzed the genetic samples of 35,027 individuals, which consist of more than 300 million variants after restricting analysis to variants without quality control flags (INFO/VFLAGS One subgroup == 0) and ABHet ratios between 0.25 and 0.75 (Table S1) [53]. The variant positions are aligned to the GRCh38 reference genome. The compressed genomic data structure (GDS) files occupy more than 60 GB of disk space. We considered only biallelic variants with minor allele counts greater than 10 and missingness rate less than 0.05. To phase the initially unphased data, identify close relatives, and determine cohorts with similar genetic ancestry, we developed an automated and reproducible bioinformatics pipeline for haplotype phasing, relatedness inference, and local ancestry inference. The pipeline is run with a single terminal command and parameter specifications in a configuration file [55]. Table S2 provides the recommended parameter settings that we use. We used the deCODE pedigree-based recombination map from 2019 [56].

#### 2.5.1 Haplotype phasing

We used `Beagle 5.4` to perform haplotype phasing [57, 58]. We used the 3,202 samples from the 1000 Genomes [59] and Human Genome Diversity Panel (HGDP) [60] data (build 38) as reference panels. We phased the roughly 20 million biallelic variants that are shared between the two datasets (Table S1); that is, we did not use `Beagle` imputation (Table S2). With 8 threads and 64 GB of random access memory (RAM), phasing chromosome 2 took 70 hours on an Intel 2.60 GHz CPU. The memory footprint of the 22 autosomal VCF files is 224 GB.

#### 2.5.2. Local ancestry inference

We used `flare` version 0.5.1 to perform local ancestry inference with the phased data [61]. We used a subset of 1,415 samples from 5 continental ancestry groups from the 1000 Genomes reference panel (Table S3), which we refer to as African (AFR), East Asian (EAS), European (EUR), South Asian (SAS), and American (AMR) ancestry groups. The smallest reference panel comprises 102 AMR individuals, whereas the largest reference panel consists of 447 AFR individuals. Browning et al. [61] showed that the average squared correlations between the true and inferred local ancestry dosages can be more than 0.90 with sample sizes this large. In Appendix A.1, we describe how we defined the AMR reference group as a union of some Peruvian in Lima, Peru, (PEL) and Mexican Ancestry in Los Angeles, California, (MXL) samples in the 1000 Genomes data. With 8 threads and 64 GB of RAM, inferring local ancestry for the variants on chromosome 2 took 28 hours on an Intel 2.60 GHz CPU. The memory footprint of the 22 autosomal VCF files is 320 GB. We made hard calls for the local ancestry with the highest probability (`probs=false` in Table S2) because storing ancestry dosages would result in an enormous memory footprint. `flare` averages the local ancestry calls for each individual to get global ancestry proportions [61].

#### 2.5.3. Defining ancestry-specific case and control cohorts

To keep our definition of AD relatively consistent, we defined AD status using the “AD” variable in the case/control phenotype files [53], excluding samples from the family-based, progressive supranuclear palsy (PSP), and Alzheimer’s Disease Neuroimaging Initiative (ADNI) datasets (see “Data and code availability” section). This binary AD variable includes a mix of 15,760 controls (coded as 0s) and 10,111 clinical or autopsy-confirmed cases (coded as 1s) that were sequenced.

Next, we stratify by ancestry because the null model assumes panmixia. We defined an initial European ancestry group as individuals inferred to have more than 90% global EUR ancestry proportion. After identifying some subsets descended from founder populations, we further split this group into two cohorts (see below). We also defined an initial African ancestry group as 2,526 individuals inferred to have more than 75% global AFR ancestry proportion. We used a smaller ancestry proportion threshold for the AFR group to ensure that there are enough cases to detect pairwise IBD.

Close familial relatedness could confound our analyses. We used `hap-ibd` [62] segments with default sequence data settings to estimate pairwise kinships with `IBDkin` [63] version 2.8.7.8 (Table S2). Within ancestry groups, we used the network-based approach of Temple et al. [40] to remove sets of individuals connected by kinship coefficients exceeding 0.125 (the expected kinship of a grandparent-grandchild pair). The relatedness network-based pruning step removed 67 samples when applied to the initial AFR ancestry cohort. The refined AFR ancestry cohort consists of 731 cases and 1,757 controls, including 1,757 females and 721 males. The mean, median, and maximum AFR ancestry proportions of the samples in the cohort are 0.84, 0.84, and 0.91, respectively. Thus, every sample in the AFR cohort has a nontrivial amount of admixture.

We identified a subset of samples from the EUR ancestry cohort with more than 10,000 IBD segments per sample; meanwhile, the mode and median of IBD segments per sample are roughly 5,000. Many samples with over 10,000 IBD segments come from the Amish Protective Variant Study, but there are more than 1000 samples with over 10,000 IBD segments that do not come from the Amish Protective Variant Study. We suspect that the remaining samples with excess IBD sharing are genetically similar to individuals from other founder populations, e.g., Ashkenazi Jewish individuals.

We analyzed the Amish and EUR ancestry samples with fewer than 10,000 IBD segments separately. We applied the relatedness network-based pruning approach to the non-Amish EUR ancestry group but not to the Amish group. The final non-Amish EUR ancestry cohort (henceforth referred to as the EUR cohort) comprises 4,783 cases and 2,841 controls, consisting of 4,502 females and 3,122 males. The mean, median, and maximum EUR ancestry proportions of the samples in the cohort are 0.99, 0.99, and 1.00, respectively. The Amish cohort comprises 95 cases and 618 controls, consisting of 437 females and 276 males.

Due to small sample sizes, we did not analyze predominant inferred EAS, SAS, or AMR ancestry cohorts in the extended case-control study. Only 55 samples have more than 66% inferred EAS ancestry. While there are 2,350 samples with more than 66% inferred SAS ancestry, there are only 22 cases. Similarly, there are 1038 individuals with more than 66% AMR ancestry but only 183 cases.

#### 2.5.4. Case-control scan

We implemented the case-control scan within the `isweep` suite of methods [40, 28]. The selection scan in `isweep` detects IBD segments in an automated and reproducible bioinformatics pipeline [55], and the case-control scan counts *post hoc* the detected IBD segments in terms of case-case and control-control pairs. We used the ≥ 2.0 cM threshold to detect IBD segments, step sizes of 0.05 cM, and a family-wise error level of 0.05 (Table S4). For the AFR and EUR analyses, we used `hap-ibd` [62] and `ibd-ends` [39] for segment detection as in Temple et al. [40] and Temple and Browning [28]. In the Amish samples, we used `hap-ibd` only for segment detection after observing issues running `ibd-ends`. (The accuracy of `ibd-ends` can be sensitive to different demographic scenarios [28].) The selection and case-control scans were run with one terminal command and configuration files (Table S4). With 112 threads and 256 GB of RAM, running `ibd-ends` on the EUR ancestry data of chromosome 2 took 2.5 hours on an Intel 2.20 GHz CPU.

After the scan, we searched the USCS Genome Browser for protein-coding genes within the significant regions. We searched the GWAS catalog to identify known associations between genetic variants and dementia or memory-related phenotypes [64, 65, 66], and we searched Agora to identify transcriptomic or proteomic evidence that a gene is associated with AD [67]. Below, we note region-specific genes whose functions are relevant to the nervous system.

## 3. Results

### 3.1 Simulation study

#### 3.1.1. Estimating the exponential decay parameter

Figure S1A shows estimates 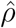 of the cross-correlation parameter from whole genome sequence data for each demographic scenario. The medians of estimates 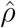 are less than 0.25 and approximately zero in the three stages of exponential growth and population bottleneck scenarios, respectively. The estimates 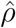 are often positive, albeit the IBD rate processes can be very noisy. Figure S1B shows that estimates 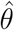 from the case-control and selection scans are highly correlated (Pearson correlation coefficient 0.48 and positive slope 0.997). Based on these estimates of 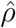 and 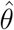, we computed the family-wise error rates of true two-dimensional OU processes of size equal to our whole genome simulations using the true *ρ* and *θ* = *θ*_1_ = *θ*_0_. Varying *ρ* did not affect the FWER. Consistent with Temple [27] and Temple and Browning [28], the discrete-spacing analytical threshold was slightly conservative when *θ* ≥ 50 and moderately conservative when *θ* ≤ 25.

#### 3.1.2. Family-wise error rate control

Table 1 shows the FWERs of the ≥ 2.0 and ≥ 3.0 cM IBD rate difference scans every 0.02 cM. Overall, the FWERs are less than 0.035, which is conservative compared to the desired rate of 0.05. Two sources of the conservativeness could be that the upper tails of the IBD rate differences are lighter than those of Gaussian distributions [35] and that the OU approximation method is conservative [38, 27]. Nevertheless, the genome-wide significance levels are greater than 5.0 *×* 10^−6^, whereas the Bonferroni-adjusted significance level is 1.0 *×* 10^−6^ (50,000 tests). The choice of segment detection threshold did not noticeably affect the FWERs, whereas ≥ 2.0 cM IBD rates can lead to anticonservative behavior in the selection scan [28]. We recommend using ≥ 2.0 cM IBD rate differences because the ≥ 3.0 cM detection threshold requires substantially more case and control samples.

**Table 1.** Genome-wide significance levels and family-wise error rates after multiple-testing corrections. Family-wise significance levels of 0.05 are adjusted for multiple testing based on scans over 10 chromosomes of size 100 cM and tests every 0.02 cM (50,000 total tests). The genome-wide significance thresholds are derived using the discrete-spacing analytical approach. The family-wise error rate (FWER) is the percentage of five hundred genome-wide scans that yield at least one statistically significant result. The demographic scenarios considered include a population bottleneck, three phases of exponential growth, and a constant population of 50,000. IBD segment detection thresholds of ≥ 2.0 and ≥ 3.0 cM are considered.

| **Demography** | **cM** | **Significance** | **FWER** |
| --- | --- | --- | --- |
| Bottleneck | 2.0 | 6.31e-6 | 0.018 |
| Staged growth | 2.0 | 5.23e-6 | 0.034 |
| Constant 50k | 2.0 | 6.66e-6 | 0.028 |
| Bottleneck | 3.0 | 9.41e-6 | 0.026 |
| Staged growth | 3.0 | 7.13e-6 | 0.030 |
| Constant 50k | 3.0 | 9.17e-6 | 0.026 |

Figure S2A-C show histograms of the standardized IBD rate difference for each demographic scenario. The OU model assumption of normally distributed IBD rate differences appears reasonable. Figure S2D-F shows the autocovariances of standardized IBD rate differences and fitted exponential decay curves. The OU model assumption of a specific autocovariance structure appears reasonable for the population bottleneck and constant population scenarios. We thus suggest that the OU model may be a good approximation for the IBD rate differences in large samples, at least to the extent that the multiple-testing thresholds control the FWER.

#### 3.1.3. Confounding due to positive selection

Figures 1 and S3 show the proportion of rejected null hypotheses when simulated directional selection is a confounding factor. For selection coefficients *s* ≥ 0.02 and sweeping allele frequencies 0.25 or 0.50, more than 15% of the time we made a Type 1 error in the ≥ 2.0 cM scan. For *s* = 0.03, nearly 50% of the time we made a Type 1 error. Recall that the hypothesis test is one-sided and that we randomly assigned case and control phenotypes; therefore, a Type 1 error rate of fifty percent is considered pathological performance. The Type 1 error rates were much smaller in the ≥ 3.0 cM scan, but sample size could be a major constraint on such scans.

**Figure 1.**
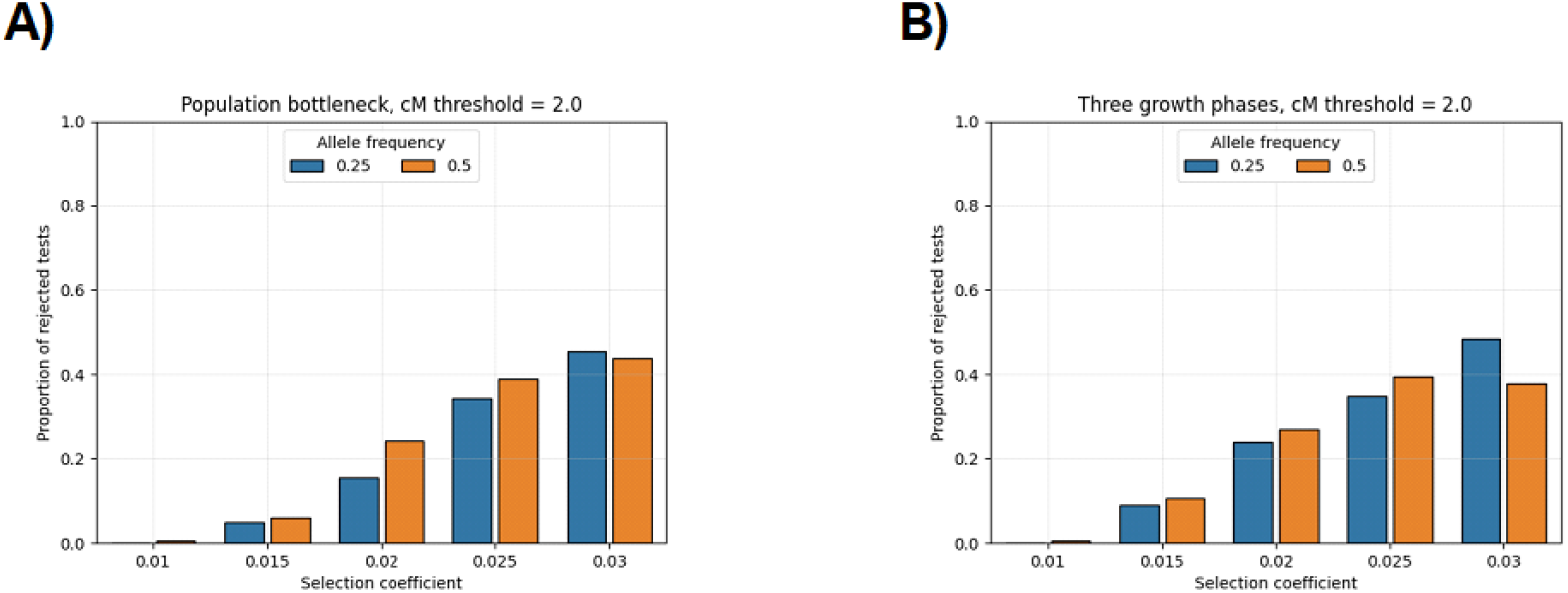
Proportion of false positives when strong positive selection is confounding. Bar plots show the proportion of times that we reject the null hypothesis of the IBD rate difference scan in terms of the selection coefficient (x-axis) and the sweeping allele frequency (colors in legend). The demographic scenarios are A) population bottleneck and B) three phases of exponential growth. Each parameter combination is simulated 200 times. The significance threshold is based on the average threshold over all null simulations. The IBD segment detection threshold is ≥ 2.0 cM.

Recent hard sweeps in a European ancestry cohort that were modeled by Temple et al. [40] had very strong (as defined by Schrider and Kern [68]) selection co-efficients between 0.01 and 0.04. For instance, Temple et al. [40] estimated a 95% selection coefficient confidence interval of (0.0278, 0.0373) at the *LCT* gene. These selection coefficient estimates align with the simulations where we observed a high probability of confounding. Juxtaposing the IBD-based selection scan [39, 40, 28] alongside the case-control scan can be an important check for confounding. We considered the hard sweep model as a process that could increase IBD sharing, but other phenomena, such as population structure, could also increase IBD sharing. Below, we demonstrate this validation step by analyzing the *LCT* gene in the inferred EUR ancestry samples.

### 3.2. The OU model assumptions are reasonable in human genetic data

One of our model assumptions is that IBD rates and rate differences are normally distributed. Figures S5A-C and S6A-C show that the empirical distributions of IBD rates and rate differences in the three ancestry cohorts resemble Gaussian distributions. The IBD rate distributions also resemble Gaussian distributions and are similar between cases and controls (Figure S7). Recall that the IBD rate and rate differences converge to a Gaussian distribution under large sample and population size conditions [35].

Our other model assumption is that the IBD rates and rate differences have exponentially decaying autocovariances. Figures S5D-F and S6D-F show that autocovariances in the three ancestry cohorts fit the OU model well. Figure S8 indicates that IBD rates in cases and controls have similar autocovariances.

For IBD rates in the AFR cohort, the exponential decay estimates for the whole sample, controls, and cases are 72, 73, and 69, respectively. For the IBD rates in the EUR cohort, the exponential decay estimates for the whole sample, control, and cases are 44, 48, and 45, respectively. These estimates are similar to those of Temple and Browning [28], who modelled IBD rates for the Trans-Omics for Precision Medicine (TOPMed) [69] samples, which are genetically similar to those of Africans and Europeans, respectively. For IBD rates in the Amish cohort, the exponential decay estimates for the whole sample, controls, and cases are 33, 33, and 34, respectively.

### 3.3. Selection scans replicate previous work in genetically similar cohorts

From the selection scans, Figure S4 shows the ≥ 2.0 cM IBD rates along the autosomes, the autosome-wide medians, the heuristic four standard deviations above the median thresholds, and discrete-spacing analytical and simulation-based thresholds for the AFR, EUR, and Amish ancestry cohorts. The genome-wide significance levels are 2.03 *×* 10^−6^, 2.91 *×* 10^−6^, and 3.66 *×* 10^−6^, respectively. There are no genome-wide significant loci in the Amish ancestry selection scan.

Table S5 provides annotations on the genome-wide significant loci in the selection scans. Many of the putatively selected loci in the TOPMed African ancestry group or the Black British group from the United Kingdom (UK) British Biobank [28] are also genome-wide significant in the ADSP African ancestry group, including at the *XYLT1* (OMIM: 608124), *SEMA3C* (OMIM: 602645), and hemoglobin beta (*HBB*, OMIM: 141900) genes. Many of the putatively selected loci in the TOPMed European ancestry group or the white British group from the UK Biobank [28] are also genome-wide significant in the ADSP European ancestry group, including at the *LCT* (OMIM: 603202), *OAS1-2-3* (OMIMs: 164350, 603350, 603351), and *HNF1B* (OMIM: 189907) genes and the major histocompatibility complex (*MHC*).

### 3.4. Identity-by-descent rates differ between cases and controls around genes associated with Alzheimer’s disease

From the case-control scans, Figure 2A shows the ≥ 2.0 cM standardized IBD rate differences along the autosomes, the autosome-wide median, and the discrete-spacing analytical threshold for the AFR, EUR, and Amish cohorts. For the IBD rate differences, the exponential decay estimates for the AFR, EUR, and Amish cohorts are 70, 70, and 24, respectively. These values correspond to genome-wide significance levels of 2.08 *×* 10^−6^, 2.08 *×* 10^−6^, and 4.47 *×* 10^−6^, respectively. Table 2 lists the risk loci where the standardized IBD rate differences exceed the ancestry-specific thresholds.

**Table 2.** Loci detected in case-control scans of the Alzheimer’s Disease Sequencing Project data. We report loci where standardized identity-by-descent (IBD) rate differences exceed the discrete-spacing analytical thresholds for the AFR ancestry, EUR ancestry, and Amish samples. The maximum standardized IBD rate difference and its corresponding p value under the null are given for each locus. Physical positions for the location of the maximum standardized IBD rate difference and the span of significance rate differences are shown in megabases (Mb). The sizes of the genome-wide significant regions are shown in centiMorgans (cM). Annotated genes are discussed in the main text, where ^*^ denotes loci where positive selection may be a confounding factor. The IBD segment detection threshold is 2.0 cM.

| Study | Chr | Max $\Delta Z$ | p value | Size (cM) | Position (Mb) | Genes |
| --- | --- | --- | --- | --- | --- | --- |
| AFR | 2 | 5.82 | 2.94e-9 | 1.50 | 4.10 (3.93-4.27) | *DCDC2C, COLEC11* |
|  | 5 | 5.56 | 1.35e-8 | 1.20 | 5.55 (5.27-5.77) | *ICE1, ADAMTS16* |
|  | 1 | 4.83 | 6.83e-7 | 0.15 | 166.38 (166.38-166.40) | . |
|  | 19 | 4.65 | 1.66e-6 | 0.05 | 16.71 (16.71-16.71) | *TMEM38A* |
| EUR | 2 | 17.02 | 2.92e-65 | 4.95 | 134.84 (133.61-139.05) | *LCT* * |
|  | 1 | 6.18 | 3.21e-10 | 0.80 | 206.00 (205.99-206.62) | * |
|  | 2 | 4.99 | 3.02e-7 | 0.20 | 15.58 (15.17-15.58) | *NBAS* |
|  | 6 | 4.91 | 4.55e-7 | 0.15 | 24.65 (24.58-24.75) | *MHC* * |
| Amish | 1 | 4.48 | 3.73e-6 | 0.15 | 177.78 (177.78-177.78) | *BRINP2, ASTN1* |

**Figure 2.**
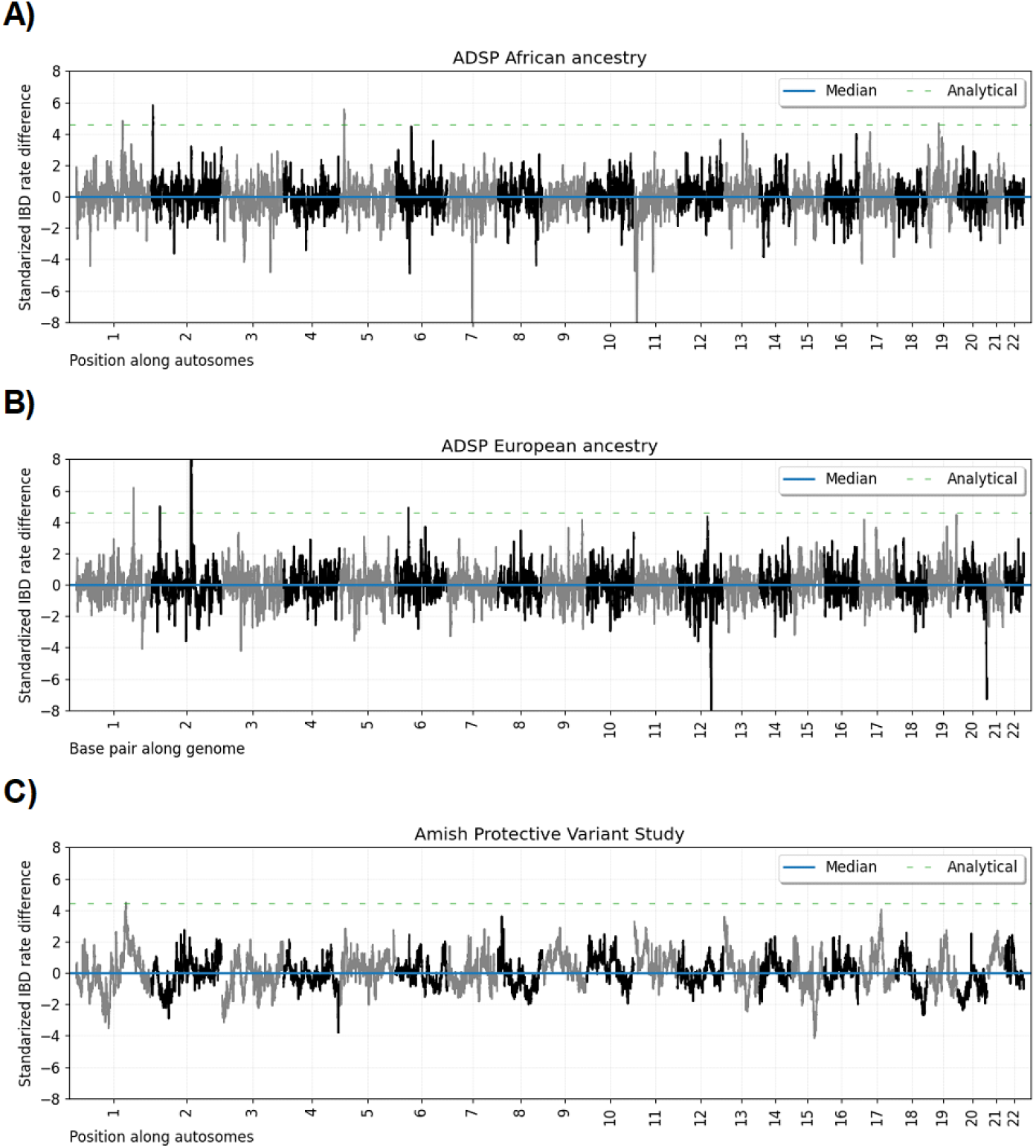
Genome-wide IBD rate difference scans for sample sets in the Alzheimer’s Disease Sequencing Project. Line plots show standardized IBD rate differences every 0.05 cM (y-axis) for base pair positions along twenty-two human autosomes. The data for each subplot is based on A) AFR ancestry, B) EUR ancestry, and C) Amish sample sets. Horizontal dashed lines show (blue) the autosome-wide median standardized IBD rate difference, (orange) the heuristic threshold of four standard deviations above the median, (green) the discrete-spacing analytical threshold, and (red) the simulation-based threshold. The IBD segment detection threshold is ≥ 2.0 cM.

Table S6 provides annotations for genes in our risk loci that are known to be associated with AD. We took these annotations of risk scores and differential expression results from the Agora web resource [67]. We considered the multi-omic risk scores developed by Cary et al. [70], differential RNA expression measured by [71], and differential protein expression in post-mortem AD individuals from Johnson et al. [72]. Three of the genome-wide significant genes in our scans have been nominated as therapeutic targets by researchers from the National Institute on Aging’s Accelerating Medicines Partnership in Alzheimer’s Disease (AMP-AD) consortium and the Target Enablement to Accelerate Therapy Development for Alzheimer’s Disease (TREAT-AD) centers.

### 3.5. Signals in the European ancestry cohort are confounded by recent selection

Figure 3 shows the significant EUR ancestry loci of the selection and case-control scans side-by-side. We suspect that many of the EUR ancestry risk loci are confounded by strong positive selection, which is indicated by asterisks in Table 2. For instance, Figure S9 shows that the excess IBD rates around the *LCT* gene have very heavy, non-Gaussian tails in both cases and controls. This situation violates the null model, resulting in IBD rate differences with heavy tails.

**Figure 3.**
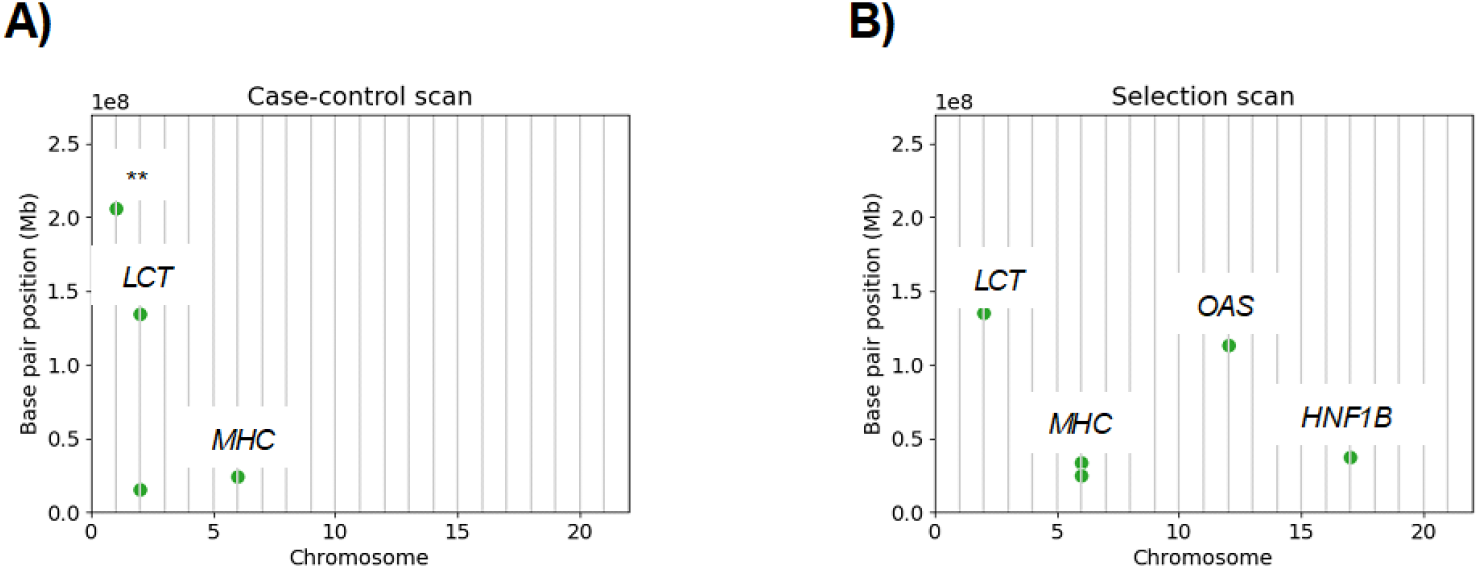
Positive selection confounds the case-control scan in European ancestry samples. The base pair position (y-axis) by chromosome number (x-axis) of genome-wide significant loci in the A) case-control and B) selection scans are shown for the European ancestry samples. Genes in the loci appearing in both A) and B) are annotated. ** This locus is identified in the TOPMed European ancestry selection scan of Temple et al. [40].

To verify that the associations between AD and *LCT* and *MHC* are spurious, we randomly assigned half of the samples to be cases and half to be controls. We reran the case-control scan with these fake phenotype labels. Figure S10 shows the ≥ 2.0 cM standardized IBD rate differences along the autosomes and the analytical significance threshold. Indeed, the *LCT* and *MHC* signal remains genome-wide significant despite the randomized phenotypes. For this randomization experiment, we used the identity-by-descent segments already inferred in the selection scan, which is the most computationally intensive step in the pipeline. Randomizing the phenotypes and rerunning the case-control scan could thus serve as a quick step to check for evidence of confounding due to selection or other mechanisms, such as population structure, that increase IBD sharing.

### 3.6. Weak associations with Alzheimer’s disease have been reported for the African-ancestry-specific risk loci

Figures 4A-B and S11 show the standardized IBD rate differences for the AFR risk loci. The strongest signal on chromosome band 2p25.3 contains no genes but lies a couple of 100 kilobase pairs (kb) from the *DCDC2C* and *COLEC11* (Ensembl: ENSG00000214866, OMIM: 612502) genes. The second strongest signal on chromosome band 5p15.32 contains the *ICE1* and *ADAMTS6* (OMIM: 617958, 605008) genes. None of the genes on chromosome band 1q24.1, the third most significant locus, have been previously associated with dementias or memory decline. The fourth significant locus on chromosome band 19p13.11 is flanked within 20 kb by the *TMEM38A* gene (OMIM: 611235).

**Figure 4.**
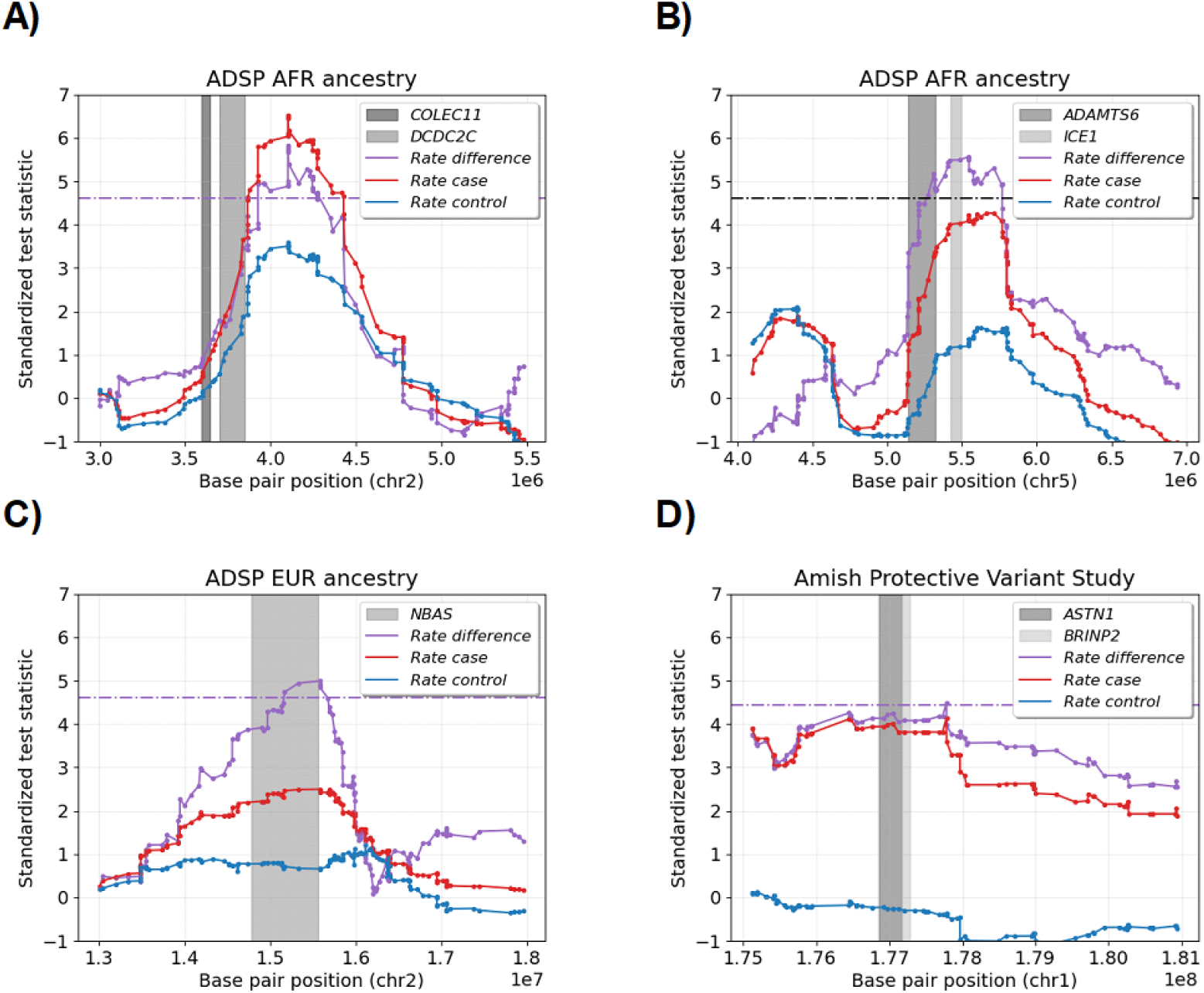
Alzheimer’s disease risk loci that are genome-wide significant in whole genome case-control scan. The scatter plot shows the standardized test statistics (y-axis) by autosomal base pair position (x-axis) for four genome-wide significant loci. The test statistics are the IBD rate difference (purple), the IBD rate in cases (red), and the IBD rate in controls (blue). The horizontal purple lines are the genome-wide significance threshold in the case-control scans. The *DCDC2C, COLEC11, ADAMTS6, ICE1, NBAS, ASTN1*, and *BRINP2* genes are shown in shades of gray. The subplot titles give the ancestry cohort.

Each of these genes has been implicated in AD, although none have yet been nominated as therapeutic targets [67]. From transcriptomic studies contrasting AD cases and controls, *COLEC11, ICE1*, and *ADAMTS6* were differentially expressed in one brain region [71], and *TMEM38A* was differentially expressed in six brain regions [71]. SNP-by-SNP interactions with *DCDC2C* and *ICE1* variants have been weakly associated with tau-related pathologies but do not reach genome-wide significance [51]. Increased expression of a dominantly inherited *ICE1* variant resulted in reduced apoptosis of neuron cells in mice [73]. Lastly, some *TMEM38A* variants have been weakly associated with sex-specific effects on memory decline but are not genome-wide significant [74].

### 3.7. Alzheimer’s disease therapeutic target NBAS is genome-wide significant in European ancestry cohort

Figure S11C shows the standardized IBD rate differences for the one non-confounded risk locus in the EUR cohort that does not appear in our selection scan (Figure 3) nor the Temple and Browning [28] scan over samples genetically similar to Europeans. The neuroblastoma-amplified sequence (*NBAS*) gene (OMIM: 608025) on chromosome band 2p24.3 is more than 600 kb long and covers most of our signal. This gene has been predicted to reduce AD progression via agonism with the *MT-ND3* mitochondrial gene (OMIM: 516002) [75, 67], which contains a known neurodegenerative protective variant (chrM:10398A*<*G) [76]. Furthermore, Agora reports high AD risk scores for the gene [67], and its RNA transcripts were differentially expressed between AD cases and controls in four brain regions [71].

### 3.8. Alzheimer’s disease therapeutic targets ASTN1 and BRINP2 are genome-wide significant in the Amish Protective Variant Study

Figure 4C shows the standardized IBD rate differences for the one genome-wide significant locus in the Amish cohort. The chromosome band 1q25.2 locus is roughly 500 kb upstream and downstream of the *ASTN1* (OMIM: 600904) and *BRINP2* (OMIM: 619359) genes, respectively. While the signal is barely genome-wide significant, recall that the multiple-testing correction is conservative. Unlike other risk loci, the elevated standardized IBD rate difference spans over 1 Mb.

Both the *ASTN1* and *BRINP2* have high AD multi-omic risk scores [67]. From transcriptomic studies contrasting AD cases and controls, *ASTN1* and *BRINP2* were differentially expressed in two and seven brain regions, respectively [71]. In tandem mass tagged [75] and liquid-free quantification data [77] from the post-mortem dorsolateral prefrontal cortex of more than 400 individuals, *ASTN1* proteins were significantly underexpressed in AD cases versus controls. In a cross-ancestry analysis, variants in *BRINP2* were associated with vascular dementia at a suggestive p value threshold [78].

The *ASTN* and *BRINP* gene families are involved in neuronal cell processes. The proteins encoded by *ASTN1* serve as adhesion molecules for migrating neurons in development phases [79]. The proteins encoded by *BRINP2* are predominantly and widely expressed in neurons and may serve a regulatory purpose [80]. Based on single cell and single nucleus transcriptomic analyses of microglia subtypes, both *ASTN1* and *BRINP2* are believed to be downregulated in AD [81].

## 4. Discussion

Here, we present an IBD-based method to detect genetic associations with binary phenotypes. The method performs a simple hypothesis test for the difference between case-case and control-control proportions of IBD sharing. Then, we determine a multiple-testing correction that adapts to dataset-specific correlations by modeling IBD rates as a stochastic process. One limitation of our approach is that we cannot directly account for population structure and admixture, but instead select ancestry-specific cohorts with reduced admixture compared to the broader consortia data. Alongside GWAS, our method could be a helpful tool to extract further genetic insights from the vast consortia data for many complex traits.

By coupling the IBD rate scan [39, 40, 28] with the IBD rate difference scan, we scrutinized the IBD mapping results with the selection results to avoid spurious inference. For instance, positive selection is known and has been demonstrated in this work to confound case-control testing in samples genetically similar to European ancestry populations. Grinde et al. [82] enumerate a list of genomic regions where selection, difficulties with sequencing and alignment, and other factors can affect human genetics analyses, knowing such genomic regions beforehand can be limiting in non-human studies. We performed scans with randomized phenotypes to scrutinize if genome-wide significant loci are confounded by other mechanisms like population structure that inflate IBD rates.

To promote ease of use, we created automated workflows to phase and perform local ancestry inference for more than 35,000 samples and more than 13,000,000 (non-ultra-rare) variants and run the case-control scan. Both pipelines are run with one command line after modifying a configuration file. The most computationally intensive step in our case-control scan is detecting IBD segments [40], which need only be run once and can be used for multiple analyses. The computational cost of IBD detection is not unreasonable, even at a biobank scale. Temple and Browning reported that IBD detection from the SNP array data of more than 400,000 samples ran in less than a week using 16 Intel 2.60 GHz CPUs.

Our scalable, interpretable, and easy-to-run approach enables systematic scans of many binary traits in hundreds of thousands of individuals. Large human biobanks with rich phenotype data [83] invite association testing of the same genotype data across multiple phenotypes. Compared to variance component methods for IBD mapping, which incur considerable costs from matrix computations [7, 8], our sample mean calculations are much faster. We doubt that extending the Cai and Browning [7] quantitative traits method to multiple binary traits will scale well to hundreds of thousands of samples.

For three distinct ancestry cohorts, we performed whole-genome scans of the IBD rates in AD cases and controls. We detected six potential genome-wide significant risk loci: four in the inferred AFR cohort, one in the inferred EUR cohort, and one in the Amish Protective Variant Study. While all six potential risk loci contain genes differentially expressed in AD individuals, none have reached genome-wide significance in late-onset AD studies. One caveat is that our approach is indirectly scanning for haplotype differences as opposed to directly testing variants, and therefore, the genes we have reported on may not be driving the signals.

Protective variants for AD [24] are of particular interest for pharmaceutical development. The one-sided IBD rate difference scan is designed to detect variants associated with a phenotype that is at low frequency in the general population but preferentially selected for in the study cohort. We analyzed samples for which a disease phenotype was ascertained, and thus we focused on searching for risk haplotypes rather than haplotypes carrying protective variants. In Appendix A.3, we extend the one-sided hypothesis test to a two-sided hypothesis test. The assumptions of the two-sided test may be reasonable when the samples are not ascertained for a specific phenotype (e.g., the UK Biobank [83]). Still, we remain skeptical about the applicability of this approach in case-control studies like ours.

Future work is required to identify specific AD risk variants driving the IBD mapping signals at genome-wide significant loci. The IBD rate difference scan narrows the signal to regions of 0.05 to 1.50 cM (Table 2). The phased variants have more than 10 minor allele counts and thus represent a tiny proportion of the sequence data (Table S1); many risk variants could be ultra-rare in the broader consortium data. One approach would be to determine if excess IBD sharing clusters [40] are enriched for cases, which could refine the search to specific haplotypes. For example, in a Colombian sample, Acosta-Uribe et al. [46] show that carriers of particular AD-related variants share overlapping IBD segments ≥ 2.0 cM on the background of African, European, and Native American local ancestry tracts.

Promisingly, some genes in the potential risk loci detected by our method have been nominated as therapeutic targets by the AMP-AD consortium. The potential risk genes *ASTN1* and *BRINP2* in the Amish cohort have been nominated as therapeutic targets, but there are no current plans for experimental validation [67]. The potential risk gene *NBAS* in the inferred EUR cohort is undergoing experimental validation as a therapeutic target [67]. On the other hand, the four potential risk loci in the AFR cohort have not been nominated as therapeutic targets. Our work inspires further investigation of these genes as therapeutic targets and ongoing biomedical research in underrepresented populations to better understand the complex genetic architecture of AD [45].

## Supporting information

Supplemental Acknowledgments

## Data and code availability

The ADSP WGS data (Accession Number NG00067) is available through qualified access. Specifically, access to the WGS data (release 4) can be requested through the National Institute on Aging Genetics of Alzheimer’s Disease (NIA-GADS) Data Sharing Site: https://dss.niagads.org/datasets/ng00067/. We used the case-control phenotypes in “ADSPCaseControlPhenotypes DS 2022.08.18.v4 ALL.csv”, which is a file available through controlled access. The columns in this phenotype file are described in the “ADSPCaseControlPhenotypes DD 2024.11.12.xls” file, which is publicly available at the NIAGADS website.

The methodology is implemented in the https://github.com/sdtemple/isweep Python package as a module, which is available under the CC0 1.0 Universal License. Phasing and local ancestry inference are performed using the workflow https://github.com/sdtemple/flare-pipeline, which has now been folded into the `isweep` package. Scripts to conduct the simulation studies are available under the v1.0 tag at https://github.com/sdtemple/isweep/papers/mult-test-paper/.

## Acknowledgments

This research has received funding from the US National Human Genome Research Institute (NHGRI) of the National Institutes of Health (NIH) under award number HG005701. S.D.T. also acknowledges funding support from the US Department of Defense National Defense Science and Engineering Graduate Fellowship (NDSEG), the US NIH T32 GM081062 Predoctoral Training Grant in Statistical Genetics (awarded to PI T.A.T.), and the Eric and Wendy Schmidt AI in Science Postdoctoral Fellowship by Schmidt Sciences, LLC. E.E.B. and N.H.C. acknowledge funding support from the US NIH and National Institute on Aging (NIA) grant 1R21AG089267. E.M.W. (co-PI), A.L.D. (co-PI), and S.H.C. acknowledge funding support from the US NIH U01 AG058589 grant.

We thank the contributors who collected samples used in this study, as well as patients and their families, whose help and participation made this work possible. Data for this study were prepared, archived, and distributed by the National Institute on Aging Alzheimer’s Disease Data Storage Site (NIAGADS) at the University of Pennsylvania (U24-AG041689), funded by the NIA. The full acknowledgment statement for the ADSP WGS data (release 4) is included supplementary material. The content of this manuscript is solely the responsibility of the authors. It does not necessarily represent the official views of the National Institutes of Health or the U.S. Department of Health and Human Services.

We thank Joshua Bis, Tyler Day, Eugene Lin, Ruoyi Cai, Sharon Browning, Elizabeth Thompson, and Tamre Cardoso for their feedback on the project. We also thank Hiep Nguyen for technical support with the cluster computing resources.

## Author contributions

S.D.T. proposed the study, planned the study, developed the method, wrote the software, and wrote the manuscript. S.D.T. and N.H.C. performed the analyses and beta tested the analysis pipeline. E.E.B. assisted with the analyses and revised the manuscript. S.H.C. and A.L.D. performed initial preprocessing and quality control of the sequence data. E.E.B., E.M.W., and T.A.T. provided advice and funding. All authors contributed to editing the manuscript.

## Consent statement

All human subjects included in this study provided informed consent.

## Web resources

- https://faculty.washington.edu/browning/beagle/b54.html: haplotype phasing (version 5.4)
- https://github.com/browning-lab/flare: local ancestry inference (version 0.5.1)
- https://github.com/browning-lab/hap-ibd: detecting identity-by-descent segments (version 1.0)
- https://github.com/browning-lab/ibd-ends: detecting identity-by-descent segments
- https://github.com/YingZhou001/IBDkin: relatedness inference (version 2.8.7.8)
- https://tskit.dev/: utilities for tree sequences, including msprime
- https://github.com/bguo068/tskibd: deriving identity-by-descent segments from tree sequences
- deCODE [56] genetic map: https://www.science.org/doi/suppl/10.1126/science.aau1043/suppl_file/aau1043_datas3.gz
- UCSC Genome Browser: https://genome.ucsc.edu
- NHGRI-EBI GWAS Catalog: https://www.ebi.ac.uk/gwas/
- Agora (accessed May 7, 2025): https://agora.adknowledgeportal.org/
- 1000 Genomes: http://ftp.1000genomes.ebi.ac.uk/vol1/ftp/data_collections/1000G_2504_high_coverage/working/20190425_NYGC_GATK/
- Human Genome Diversity Project high-coverage sequence data: ftp://ngs.sanger.ac.uk/production/hgdp/hgdp_wgs.20190516/
- gnomAD principal components (v3): https://gnomad.broadinstitute.org/downloads
- Training a random forest classifier with principal components (October 15, 2021 blog): https://gnomad.broadinstitute.org/news/

### A. Appendix

#### A.1. Determining an American ancestry reference panel

We used the gnomAD principal component variant loadings [84] to train a random forest classifier that predicts ancestry labels with the 1000 Genomes [59, 85] and HGDP [60] samples. The 61 Indigenous American ancestry samples in HGDP are Colombian (7), Karitiana (12), Maya (21), Pima (13), and Surui (8). We held out the 1000 Genomes samples with the superpopulation label AMR, which includes admixed samples. With the trained random forest, we predicted the ancestry label of the held-out samples.

Ultimately, we defined our AMR reference panel in the 1000 Genomes data as those samples with predicted local probabilities of Indigenous American ancestry exceeding 0.90. Our strategy resembles how Jiménez-Kaufmann et al. [86] augmented their Native American reference and imputation panel with admixed samples. The accuracy of `flare` improves as the size of the smallest reference panel increases [61]. In future work, we will perform local ancestry inference with more of the 1000 Genomes and HGDP samples as a reference panel than the 1415 reference samples that we used here.

#### A.2. Simulation-based approach to multiple testing

Another way to control the FWER is to simulate two-dimensional OU processes given estimates 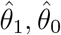, and 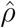. We use the Pearson correlation coefficient of 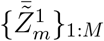 and 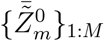 to estimate 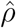. We regress the log autocovariances 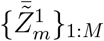 and 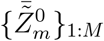 by genetic distance to estimate 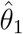 and 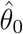, respectively [28]. Now, let the correlation matrix **P** be such that **P**_1,1_ = **P**_2,2_ = 1 and 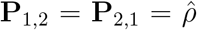, and the let **I**_2*×*2_ be the identity matrix. The lower triangular matrix of the Cholesky decomposition of **P** is denoted as **Q**.

The simulation approach follows the general procedure to simulate a multidimensional OU process. Let *J* and *M* := ⌊*L ÷ δ*⌋ be the numbers of simulations and markers, respectively, where *L* is the total length of the chromosomes analyzed, and *δ* is the step size (in Morgans). To simplify things, we simulate a single chromosome of the total genome length *L* instead of simulating the individual two-dimensional OU processes with different chromosome lengths.

##### Algorithm 1

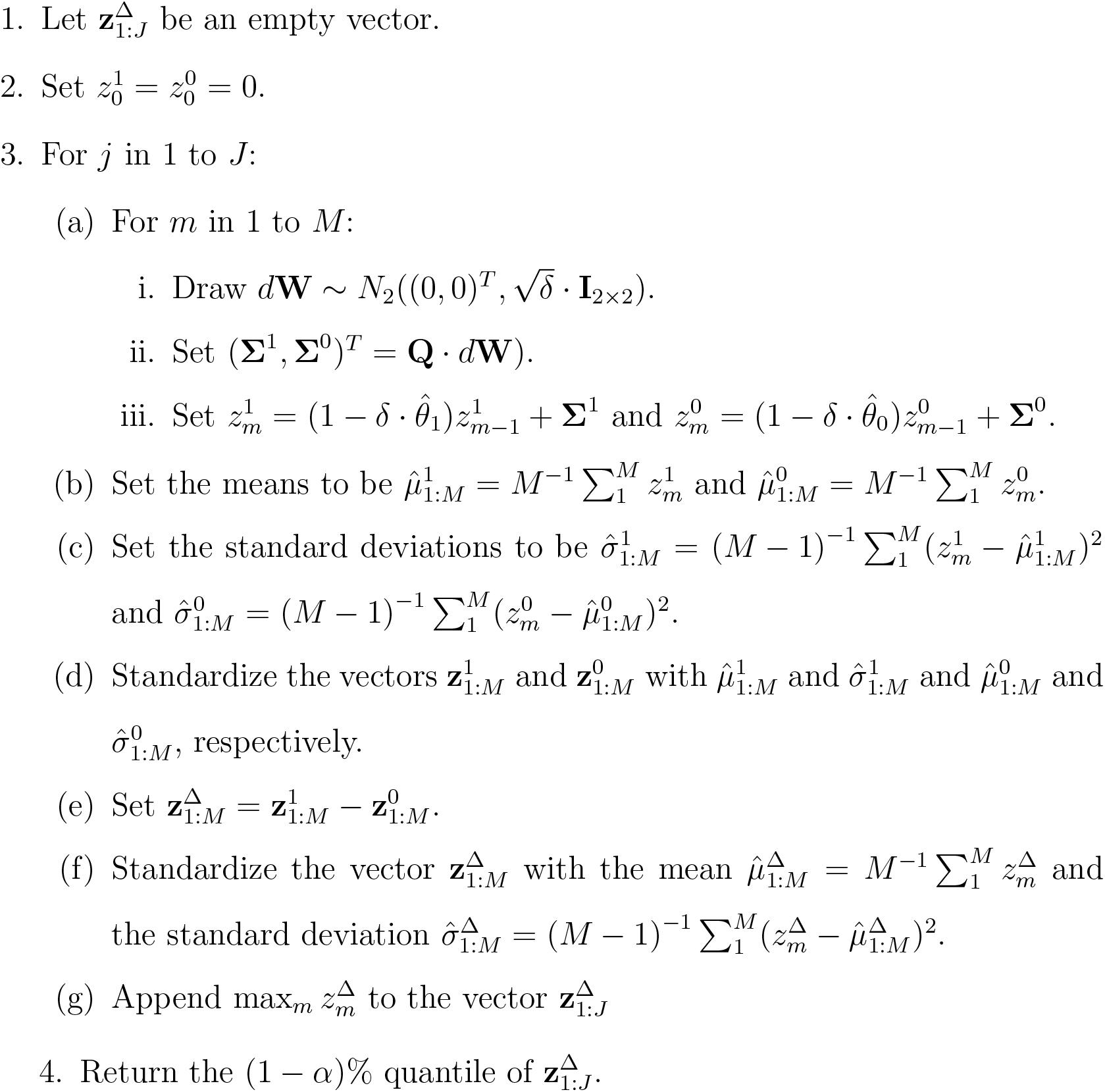

This simulation approach requires a few thousand simulations of the entire genome to calculate thresholds for family-wise significance levels like 0.05. It runs within a few minutes (depending on the total length *L* of the chromosomes) on an Intel 2.60 GHz CPU. This multiple-testing correction approximately controls the FWER when the true model is the OU process (Figure S1 and Temple [27]).

#### A.3. The two-sided test

The alternative model in the two-sided hypothesis test is 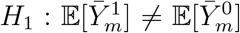 instead of 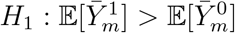, where *µ* is a genome-wide mean IBD rate around a locus. Recall that 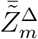 is the standardized difference of IBD rates and 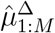 and 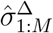 are genome-wide average and standard deviation of the standardized differences 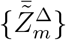. Then, the two-sided test is

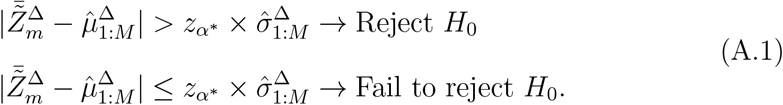

From Siegmund and Yakir [38], the multiple-testing threshold 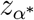 is determined by our analytical approach, except that the family-wise error rate is now

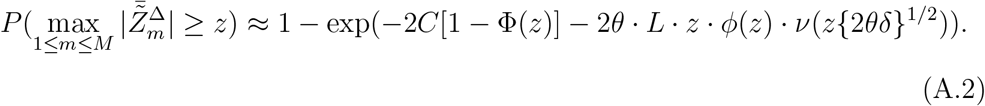

To not decrease power unnecessarily, we recommend the one-sided test unless one believes that the alternatives 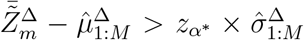 (excess IBD rates in case individuals) and 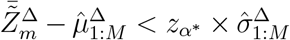 (excess IBD rates in control individuals) are equally likely. Based on the ascertainment scheme, we believe the one-sided test is more appropriate than the two-sided test for our AD study.

## Supplementary figures

**Figure S1.**
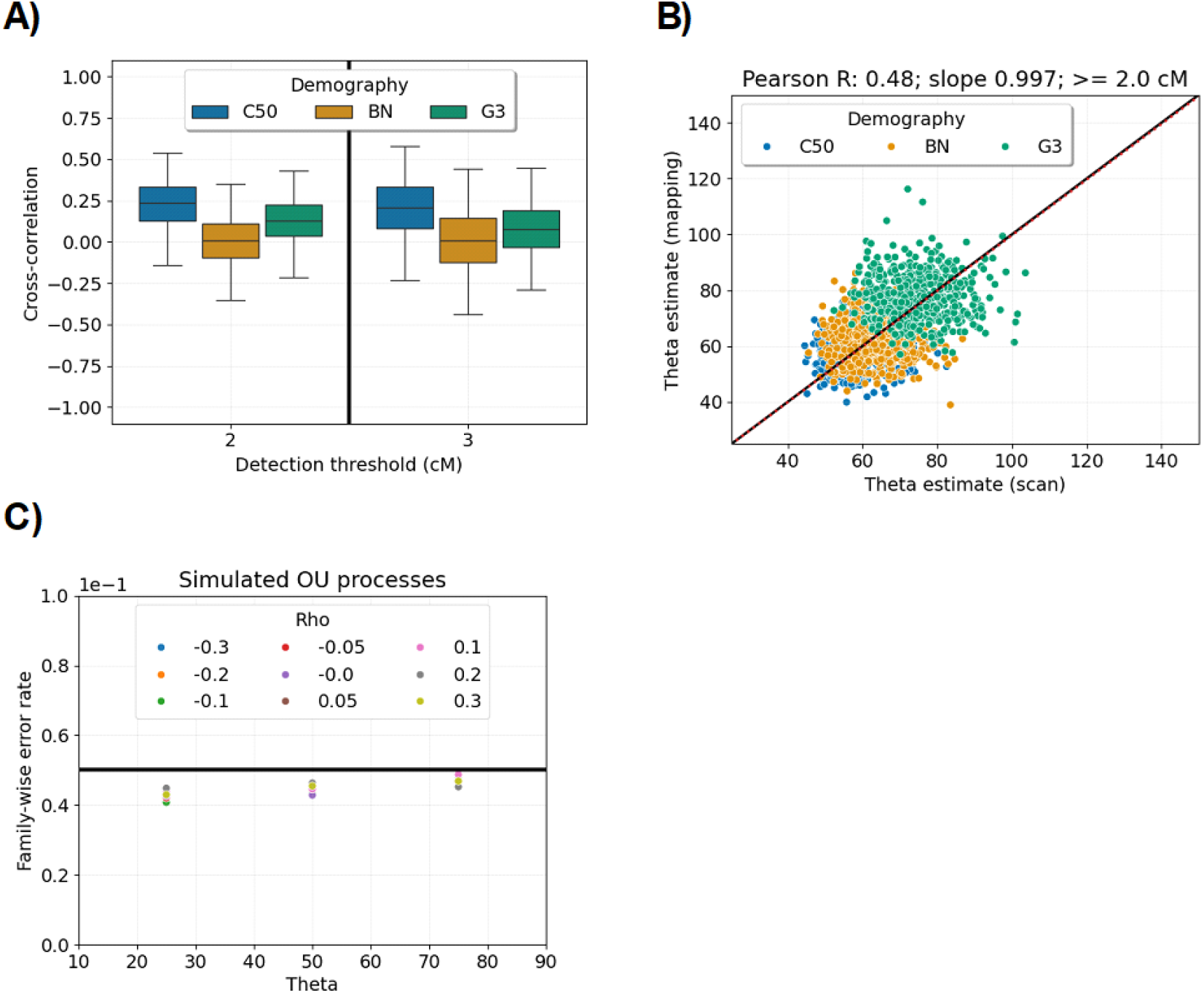
Robustness to nonzero cross-correlations. A) Boxplots show the 1st, 25th, 50th, 75th, and 99th percentiles of estimated correlations (y-axis) of case and control IBD rates with different IBD segment detection thresholds (x-axis). B) Estimates of the exponential decay parameter in the case-control scan (y-axis) are plotted versus the selection scan (x-axis). The simulated data come from a constant 50k (blue), population bottleneck (orange), and three phases of exponential growth (green) demographic scenarios. The case-control scan is performed by randomly assigning half of the 2500 samples to the case phenotype. C) The family-wise error rates (y-axis) of the standardized difference scan are plotted in terms of the exponential decay parameter *θ* (x-axis) and the cross-correlation *ρ* (legend). The desired family-wise error rate is 0.05 (indicated by the horizontal black line). The true Ornstein-Uhlenbeck process is simulated, and the true *θ* is used in calculating the discrete-spacing analytical threshold.

**Figure S2.**
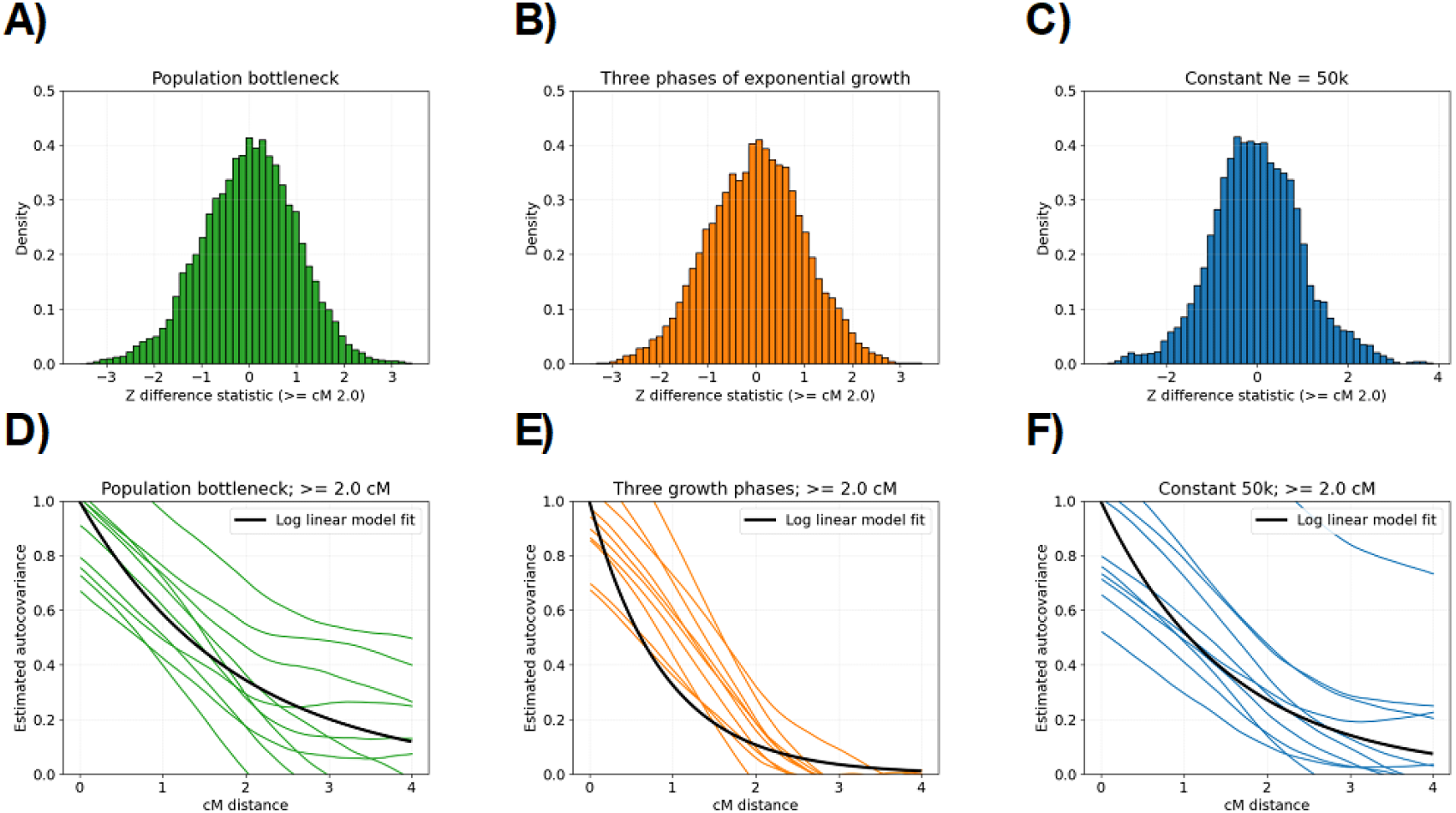
Distribution and autocovariance of standardized IBD rate differences in example data. A-C) The standardized IBD rate differences are plotted as histograms for population bottleneck (green), three phases of exponential growth (green), and constant 50k (blue) demographic simulations. Each histogram has 50 bins, and the x-axis ranges from -5 to 5. D-F) In estimating the exponential decay parameters for the IBD rate difference process, each colored line shows estimated autocovariances (y-axis) for different cM distances (x-axis) and a specific chromosome. The black lines represent the predicted autocovariances from the fitted Ornstein-Uhlenbeck processes using the estimated parameters 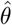. Each plot represents the results of one simulated example. The IBD segment detection threshold is ≥ 2.0 cM.

**Figure S3.**
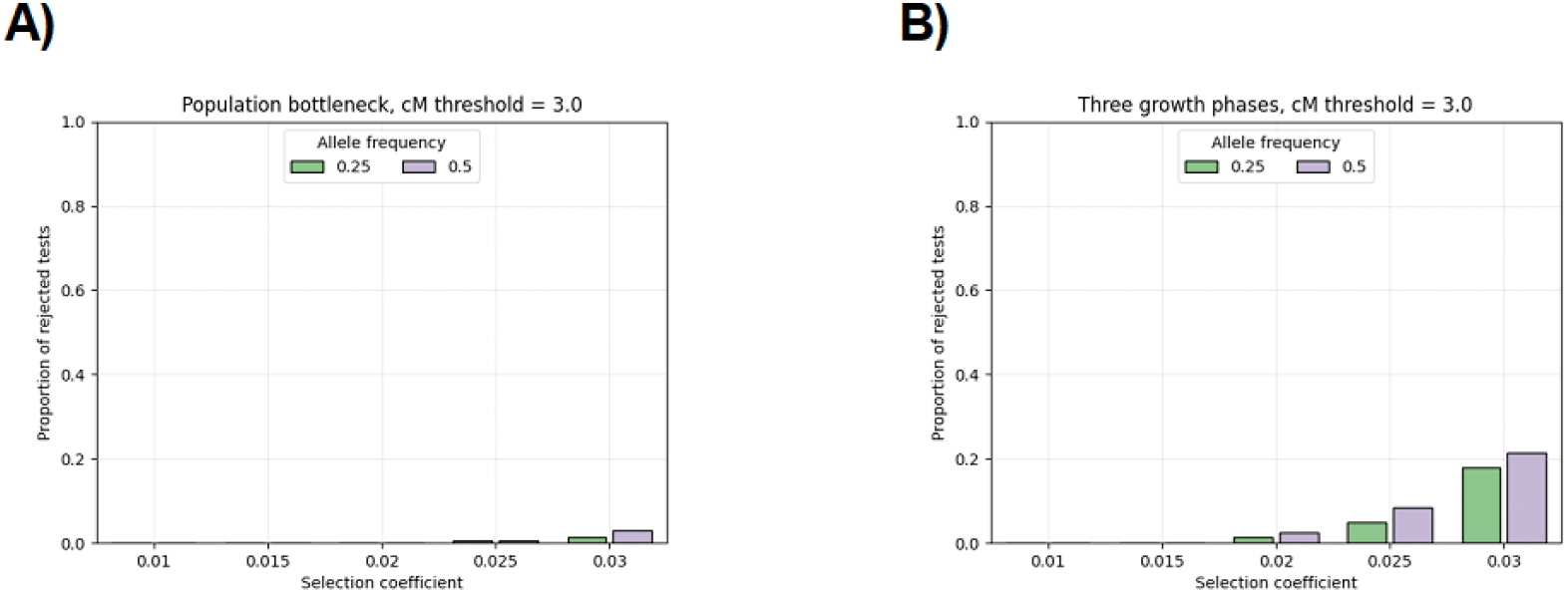
Proportion of false positives when strong positive selection is confounding and IBD segments are longer than 3.0 cM. Bar plots show the proportion of times that we reject the null hypothesis of the IBD rate difference scan in terms of the selection coefficient (x-axis) and the sweeping allele frequency (colors in legend). The demographic scenarios are A) population bottleneck and B) three phases of exponential growth. Each parameter combination is simulated 200 times. The significance threshold is based on the average threshold over all null simulations.

**Figure S4.**
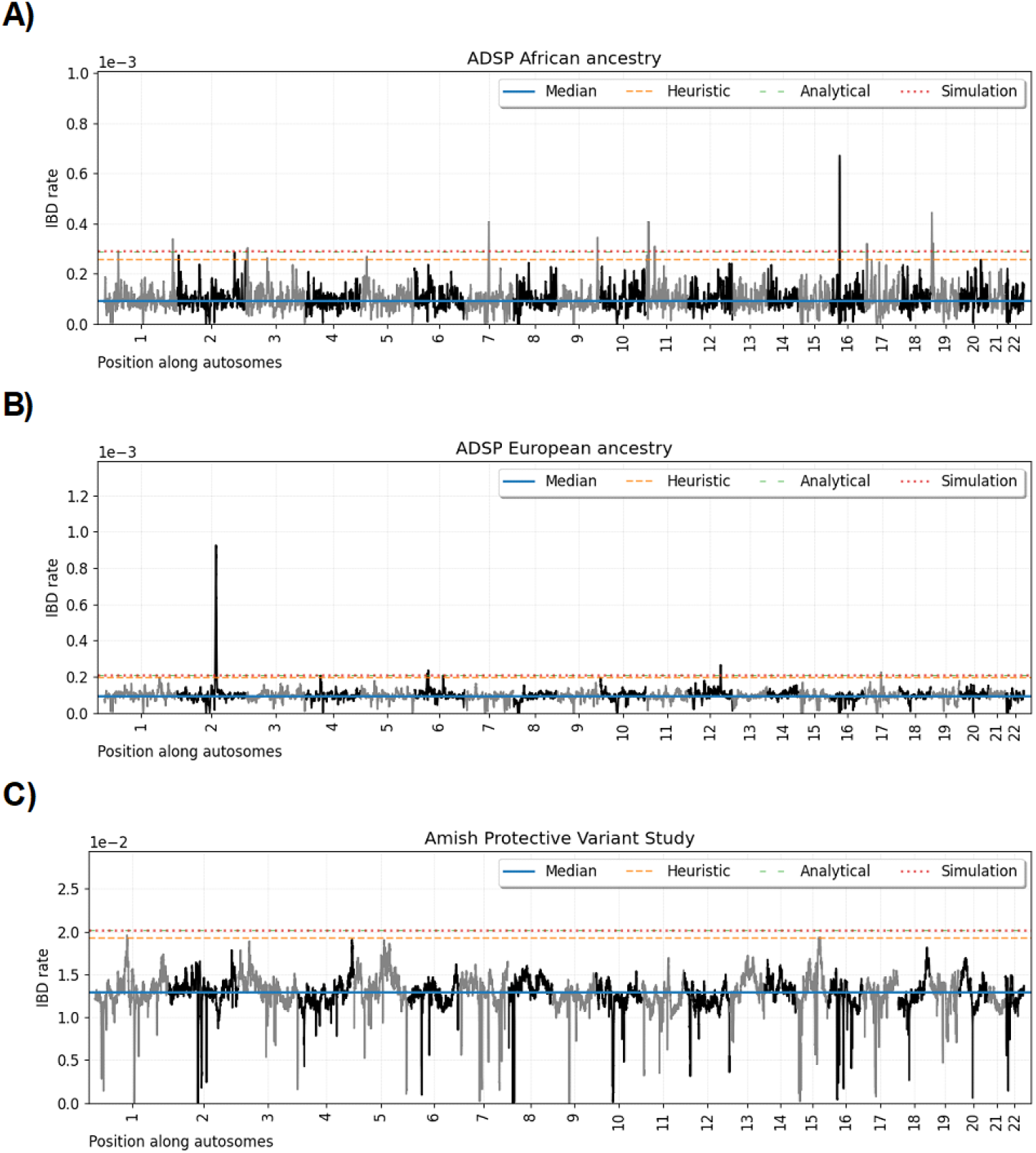
Genome-wide IBD rate scans for sample sets in the Alzheimer’s Disease Sequencing Project. Line plots show IBD rates every 0.05 cM (y-axis) for base pair positions along twenty-two human autosomes. The data for each subplot is based on A) AFR ancestry, B) EUR ancestry, and C) Amish sample sets. Horizontal dashed lines show (blue) the autosome-wide median IBD rate, (orange) the heuristic threshold of four standard deviations above the median, (green) the discrete-spacing analytical threshold, and (red) the simulated-based threshold. The analytical and simulation-based thresholds differ by less than 5e-6. The IBD segment detection threshold is ≥ 2.0 cM.

**Figure S5.**
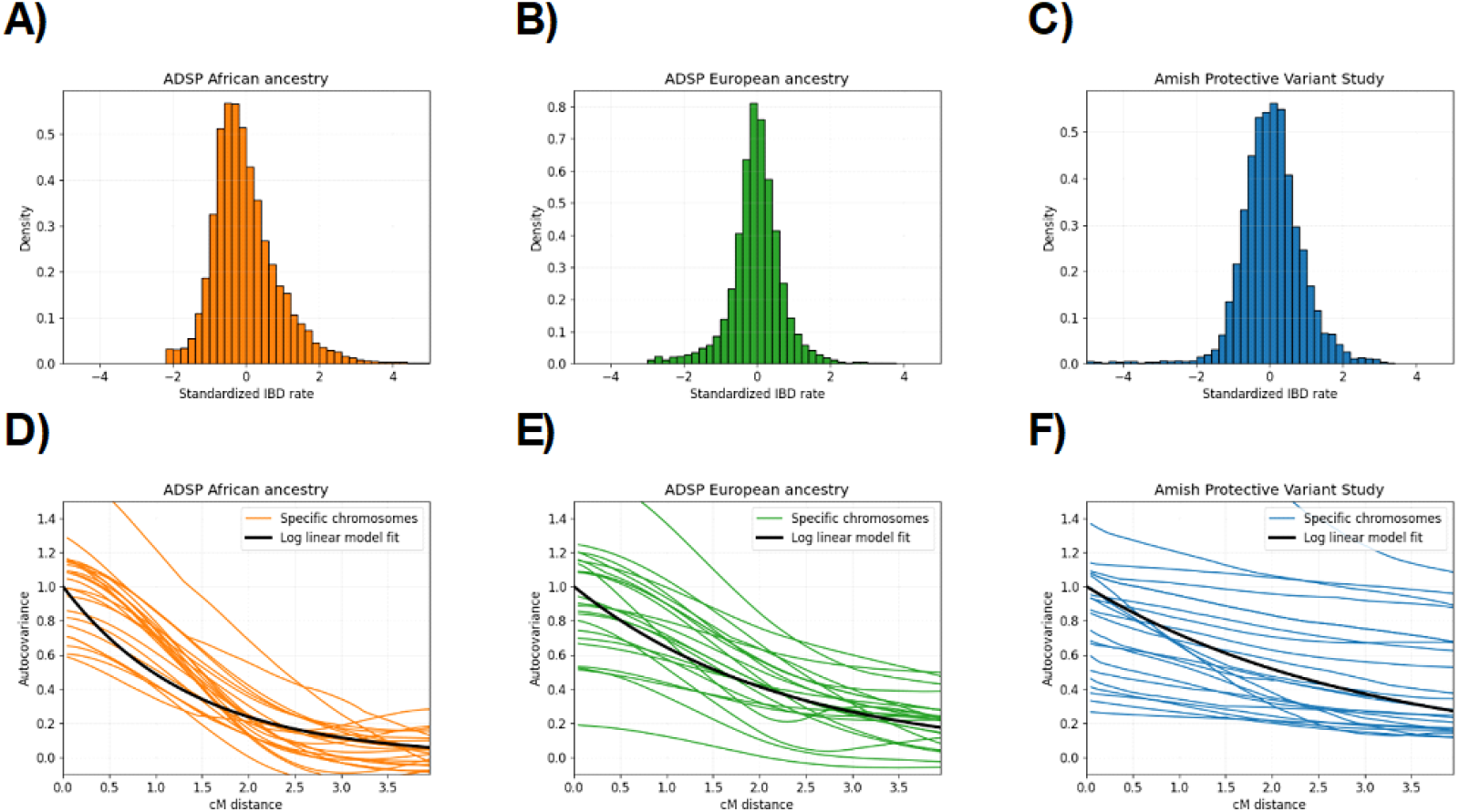
Distribution and autocovariance of IBD rates in selection scans. A-C) The standardized IBD rates are plotted as histograms for AFR ancestry (orange), EUR ancestry (green), and Amish (blue) sample sets. Each histogram has 50 bins, and the x-axis ranges from -5 to 5. D-F) In estimating the exponential decay parameters for the IBD rate process, each colored line shows estimated autocovariances (y-axis) for different cM distances (x-axis) and a specific chromosome. The black lines represent the predicted autocovariances from the fitted Ornstein-Uhlenbeck processes using the estimated parameters 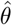. Exponential decay estimates of the AFR, EUR, and Amish ancestry samples are 72, 44, and 33.

**Figure S6.**
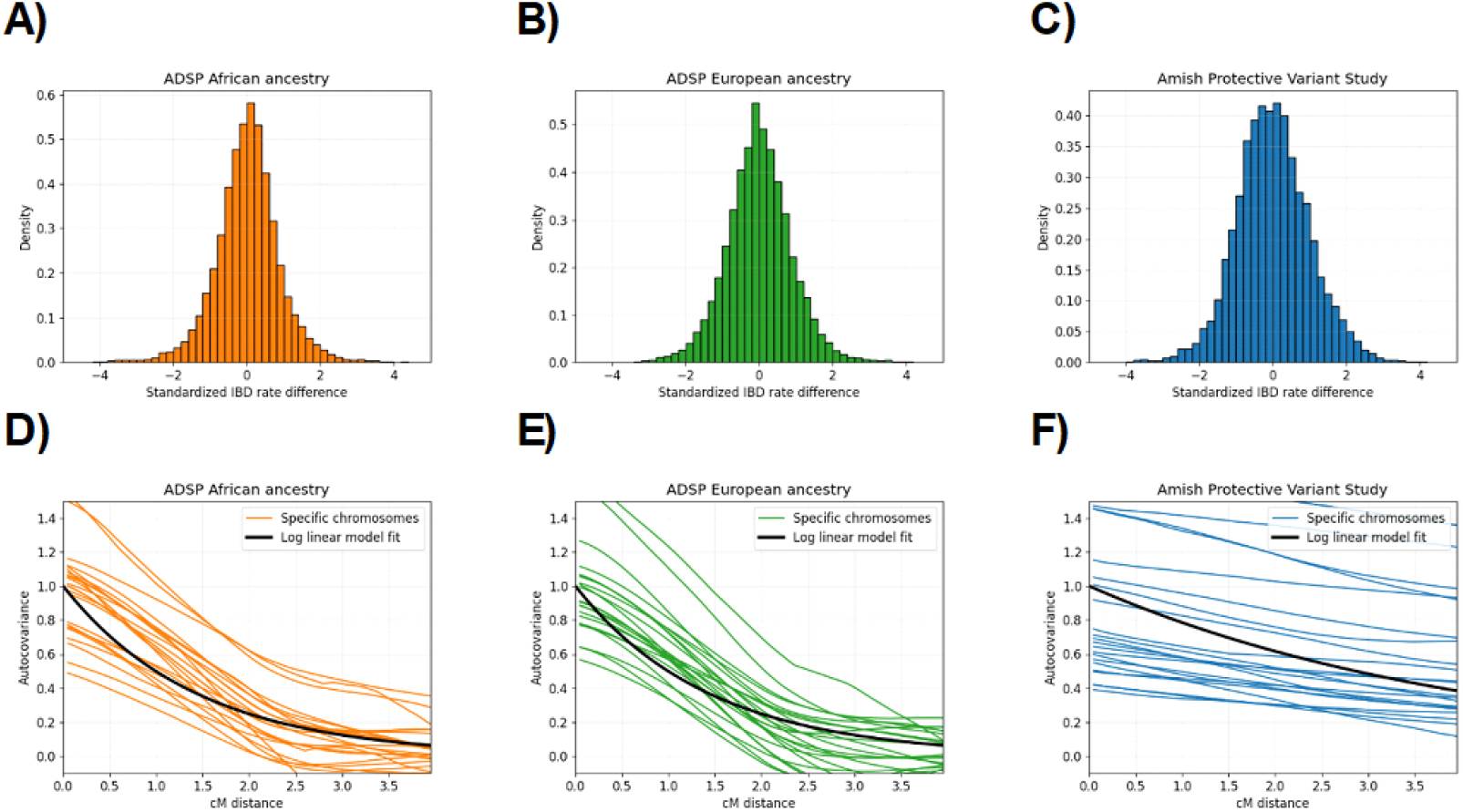
Distribution and autocovariance of standardized IBD rate differences in case-control scans. A-C) The standardized IBD rate differences are plotted as histograms for AFR ancestry (orange), EUR ancestry (green), and Amish (blue) sample sets. Each histogram has 50 bins, and the x-axis ranges from -5 to 5. D-F) In estimating the exponential decay parameters for the IBD rate difference process, each colored line shows estimated autocovariances (y-axis) for different cM distances (x-axis) and a specific chromosome. The black lines represent the predicted autocovariances from the fitted Ornstein-Uhlenbeck processes using the estimated parameters 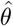. Exponential decay estimates of the AFR, EUR, and Amish ancestry control samples are 70, 70, and 24.

**Figure S7.**
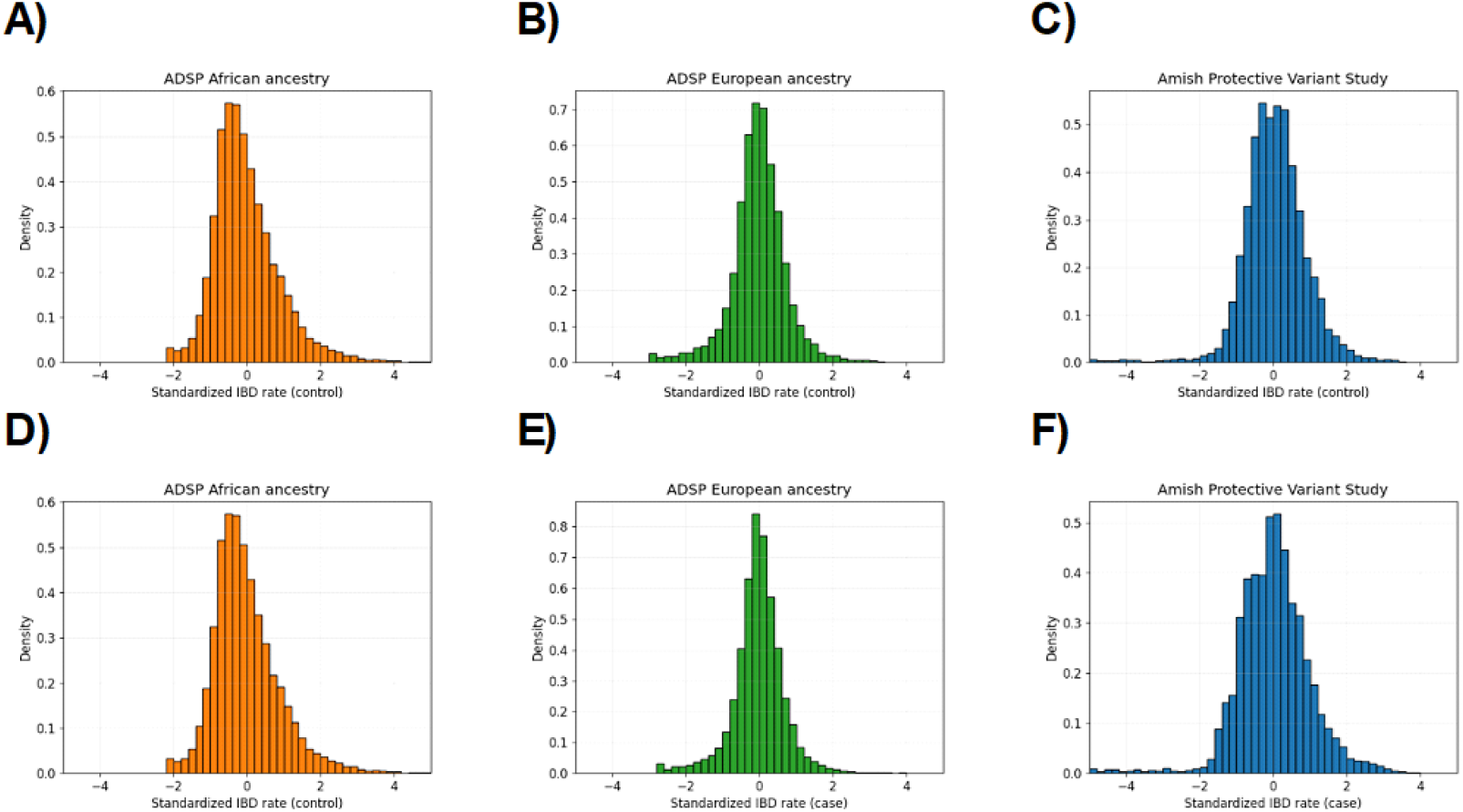
Histograms of standardized IBD rates in cases and controls. The standardized IBD rates ≥ 2.0 cM (*x*-axis) of controls A-C) and cases D-F) are shown for AFR (orange), EUR (green), and Amish (blue) sample sets. Each histogram has fifty bins, and the x-axes range from -5 to 5.

**Figure S8.**
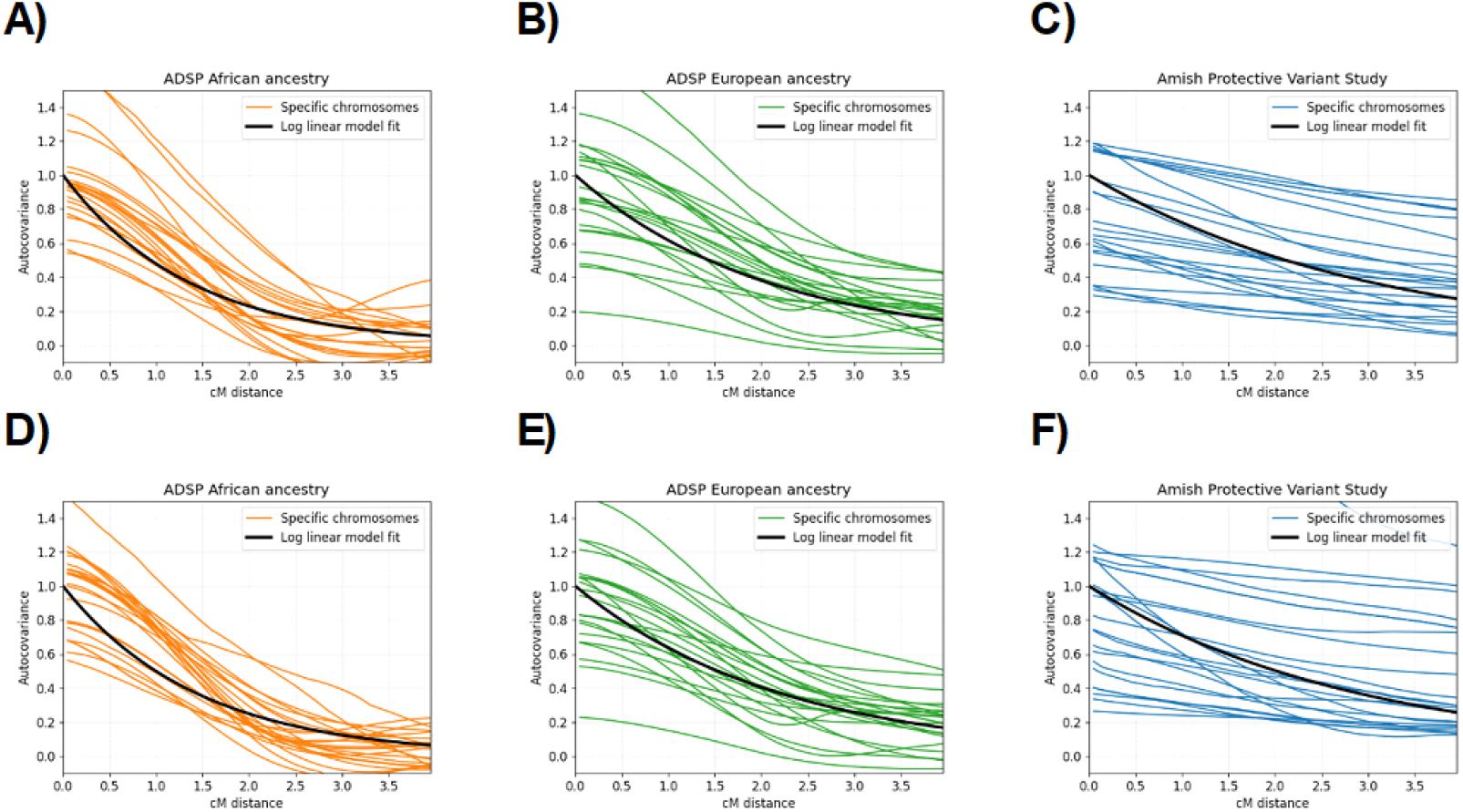
Estimating exponential decay parameter *θ* in the cases and controls separately. Each colored line shows estimated autocovariances (y-axis) for different cM distances (x-axis) and a specific chromosome. The black lines represent the predicted autocovariances from the fitted Ornstein-Uhlenbeck processes using the estimated parameters 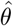. The subplots A-C) and D-F) show results for case and control sample sets, respectively. The data for each subplot is based on AFR ancestry (orange), EUR ancestry (green), and C) Amish (blue) sample sets. Exponential decay estimates for the AFR, EUR, and Amish ancestry control samples are 73, 48, and 33, respectively. Exponential decay estimates of the AFR, EUR, and Amish ancestry control samples are 69, 45, and 34.

**Figure S9.**
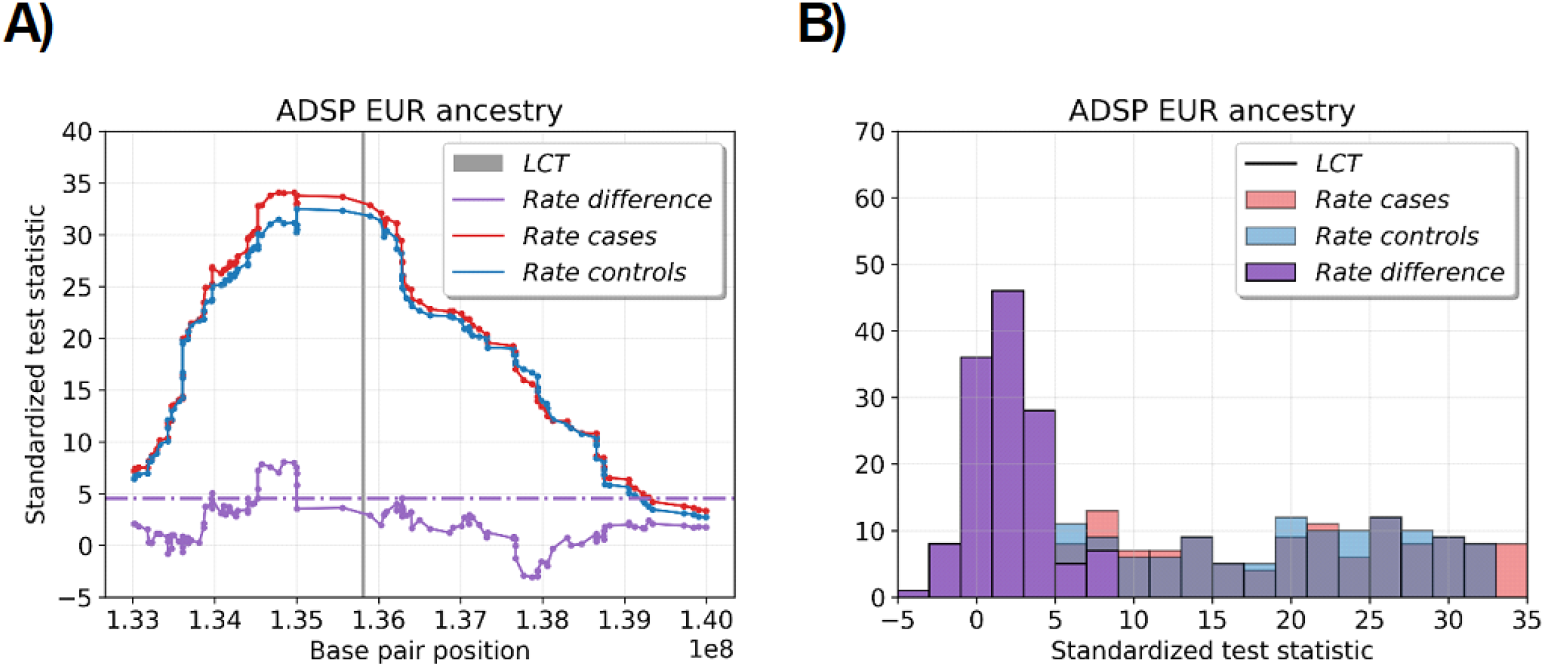
Randomized analysis of *LCT* gene. A) The line plot and B) the histogram show standardized test statistics every 0.05 cM for base pair positions around the *LCT* gene. The data is based on ≥ 2.0 cM IBD segments in all (purple), case (red), and control (blue) samples from the European ancestry cohort. We randomly assign half of the samples to be cases and the other half to be controls. A) The horizontal purple line shows analytical significance thresholds, and the gray span covers the *LCT* gene.

**Figure S10.**
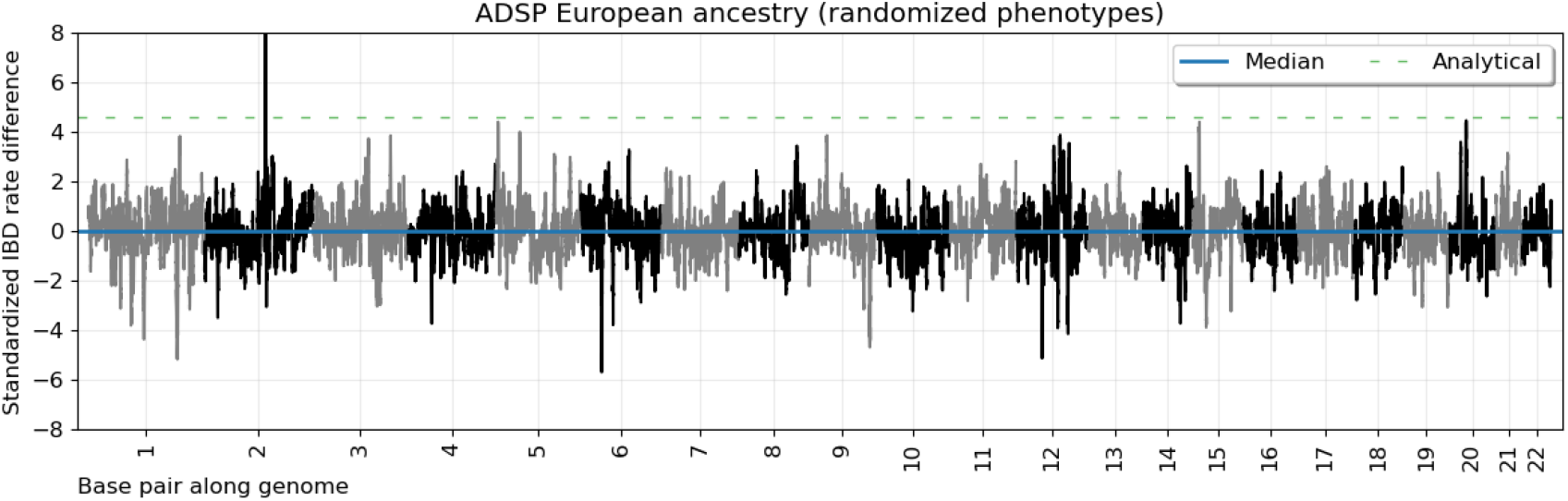
IBD rate difference scan with randomized binary phenotypes. Line plots show standardized IBD rate differences every 0.05 cM (y-axis) for base pair positions along twenty-two human autosomes. The data is based on ≥ 2.0 cM IBD segments in the EUR ancestry samples. We randomly assign half of the samples to be cases and the other half to be controls. Horizontal dashed lines show (blue) the autosome-wide median IBD rate, (orange) the heuristic threshold of four standard deviations above the median, (green) the discrete-spacing analytical threshold, and (red) the simulated-based threshold.

**Figure S11.**
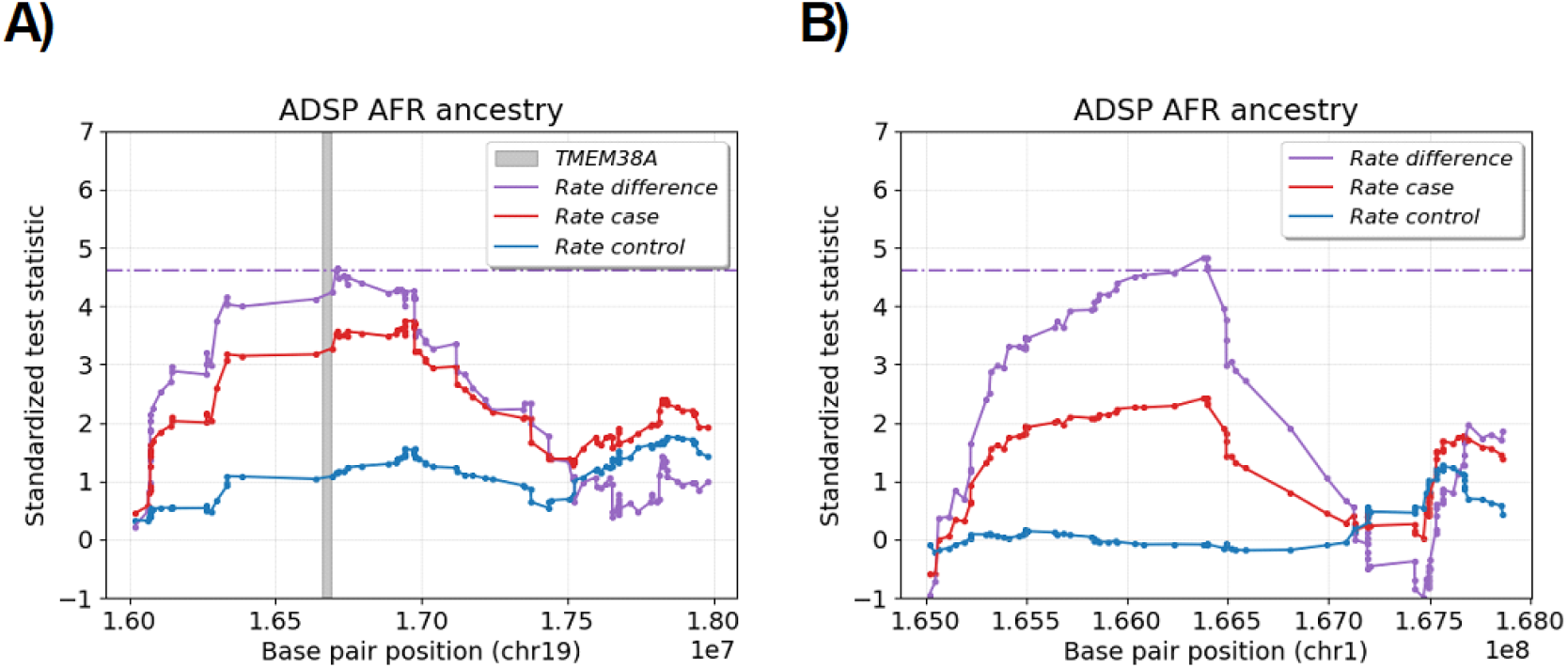
Two African ancestry-specific risk loci that are genome-wide significant in Alzheimer’s disease case-control scan. The scatter plot shows the standardized test statistics (y-axis) by autosomal base pair position (x-axis) for two loci that are genome-wide significant. The test statistics are the IBD rate difference (purple), the IBD rate in cases (red), and the IBD rate in controls (blue). The horizontal purple lines are the genome-wide significance threshold in the case-control scan. A) The *TMEM38A* gene is shown in a gray shade.

## Supplementary tables

**Table S1.**
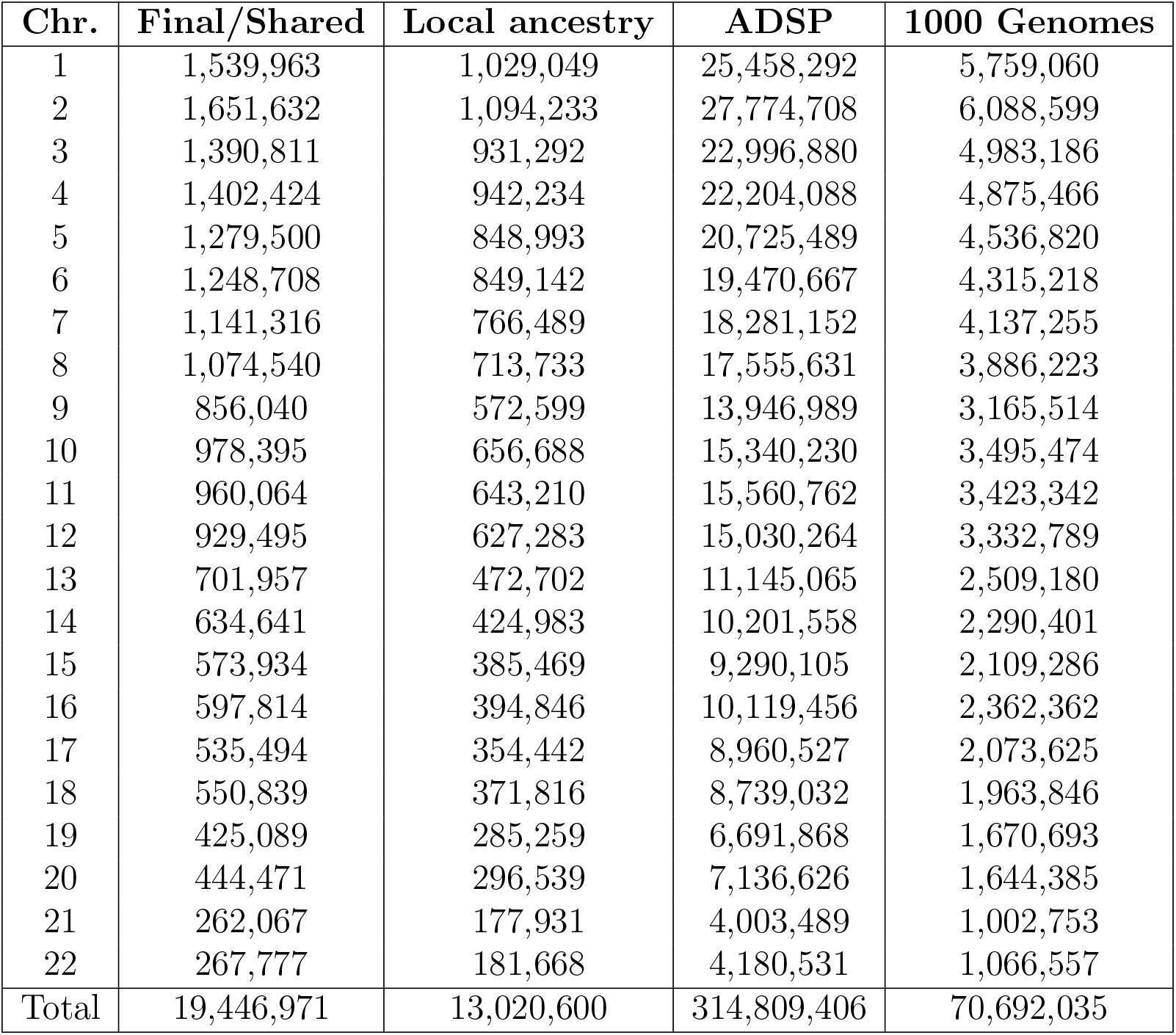
Number of phased and ancestry-inferred variants. The variant counts are given for the shared and phased variants, the variants with local ancestry dosages, the initial unphased ADSP data, and the initial phased 1000 Genomes and HGDP data.

| **Chr.** | **Final/Shared** | **Local ancestry** | **ADSP** | **1000 Genomes** |
| --- | --- | --- | --- | --- |
| 1 | 1,539,963 | 1,029,049 | 25,458,292 | 5,759,060 |
| 2 | 1,651,632 | 1,094,233 | 27,774,708 | 6,088,599 |
| 3 | 1,390,811 | 931,292 | 22,996,880 | 4,983,186 |
| 4 | 1,402,424 | 942,234 | 22,204,088 | 4,875,466 |
| 5 | 1,279,500 | 848,993 | 20,725,489 | 4,536,820 |
| 6 | 1,248,708 | 849,142 | 19,470,667 | 4,315,218 |
| 7 | 1,141,316 | 766,489 | 18,281,152 | 4,137,255 |
| 8 | 1,074,540 | 713,733 | 17,555,631 | 3,886,223 |
| 9 | 856,040 | 572,599 | 13,946,989 | 3,165,514 |
| 10 | 978,395 | 656,688 | 15,340,230 | 3,495,474 |
| 11 | 960,064 | 643,210 | 15,560,762 | 3,423,342 |
| 12 | 929,495 | 627,283 | 15,030,264 | 3,332,789 |
| 13 | 701,957 | 472,702 | 11,145,065 | 2,509,180 |
| 14 | 634,641 | 424,983 | 10,201,558 | 2,290,401 |
| 15 | 573,934 | 385,469 | 9,290,105 | 2,109,286 |
| 16 | 597,814 | 394,846 | 10,119,456 | 2,362,362 |
| 17 | 535,494 | 354,442 | 8,960,527 | 2,073,625 |
| 18 | 550,839 | 371,816 | 8,739,032 | 1,963,846 |
| 19 | 425,089 | 285,259 | 6,691,868 | 1,670,693 |
| 20 | 444,471 | 296,539 | 7,136,626 | 1,644,385 |
| 21 | 262,067 | 177,931 | 4,003,489 | 1,002,753 |
| 22 | 267,777 | 181,668 | 4,180,531 | 1,066,557 |
| **Total** | **19,446,971** | **13,020,600** | **314,809,406** | **70,692,035** |

**Table S2.** Parameter settings for phasing, local ancestry inference, and relatedness inference.

| Software | Parameter | Value |
| --- | --- | --- |
| Beagle 5.4 | impute | false |
|  | window | 5.0 |
| flare | min-maf | 0.005 |
|  | min-mac | 10 |
|  | gen | 10 |
|  | probs | false |
| hap-ibd | min-seed | 1.0 |
|  | min-extend | 0.2 |
|  | min-output | 2.0 |
|  | min-mac | 10 |
|  | max-gap | 1000 |
|  | min-markers | 100 |

**Table S3.** Sample sizes for reference panels. The abbreviations are Gambian in Western Division, The Gambia (GWD), Luhya in Webuye, Kenya (LWK), Yoruba in Ibadan, Nigeria, Han Chinese in Beijing, China (CHB), Southern Han Chinese (CHS), Japanese in Tokyo, Japan (JPT), Utah residents with Western and Northern European ancestry (CEU), British in England and Scotland (GBR), Toscani in Italy (TSI), Bengali in Bangladesh (BEB), Mexican Ancestry in Los Angeles, California (MXL), and Peruvian in Lima, Peru (PEL).

| **Superpopulation** | **Subpopulation** | **Sample size** |
| --- | --- | --- |
| African (AFR) | GWD | 176 |
|  | LWK | 97 |
|  | YRI | 174 |
|  | Total | 447 |
| East Asian (EAS) | CHB | 103 |
|  | CHS | 163 |
|  | JPT | 102 |
|  | Total | 368 |
| European (EUR) | CEU | 174 |
|  | GBR | 90 |
|  | TSI | 103 |
|  | Total | 367 |
| South Asian (SAS) | BEB | 131 |
| American (AMR) | MXL | 14 |
|  | PEL | 88 |
|  | Total | 102 |

**Table S4.** Parameter settings for selection and case-control scans. If not specified, default settings for sequence data are otherwise used in `hap-ibd` and `ibd-ends`. For the African and European ancestry analyses, we use `hap-ibd` for candidate segments and `ibd-ends` for refining the segment endpoints. For the Amish analysis, we only use `hap-ibd` with the sequence data settings. The `ibd-ends` error rate is set to 1e-4 based on initial estimates from preliminary analyses of chromosomes 20 to 22.

| Software | Parameter | Value |
| --- | --- | --- |
| **hap-ibd**<br>(EUR & AFR) | min-seed | 1.0 |
|  | min-extend | 0.2 |
|  | min-output | 2.0 |
|  | min-maf | 0.40 |
| **hap-ibd**<br>(Amish) | min-seed | 0.5 |
|  | min-extend | 0.2 |
|  | min-output | 1.0 |
|  | min-maf | 0.01 |
| **ibd-ends** | err | 1.0e-4 |
|  | quantiles | 0.5 |
|  | min-maf | 0.001 |
| Selection and<br>case-control<br>scans | step_size_cm | 0.05 |
|  | scan_cutoff | 2.0 |
|  | confidence_level | 0.05 |
|  | num_sims | 500 |
|  | cm_gap | 0.5 |
|  | outlier_cutoff | 4.0 |
|  | auto_covariance_length | 4.0 |
| Selection scan | covered_cm_region | 0.50 |

**Table S5.** Loci detected in selection scans of the Alzheimer’s Disease Sequencing Project data. We report loci where identity-by-descent (IBD) rates exceed the discrete-spacing analytical thresholds for the AFR ancestry and EUR ancestry samples. The maximum IBD rate is given for each locus. Physical positions for the location of the maximum IBD rate in loci where excess IBD rates span more than 0.5 cM are shown in megabases (Mb). The sizes of the excess IBD regions are shown in centiMorgans (cM). Annotated genes or gene complexes are discussed in the main text. The IBD segment detection threshold is 2.0 cM.

| Study | Chr | Max rate (1e-4) | Size (cM) | Position (Mb) | Genes |
| --- | --- | --- | --- | --- | --- |
| EUR | 2 | 9.25 | 6.55 | 135.00 (132.88-139.25) | *LCT* |
|  | 12 | 2.64 | 2.15 | 113.09 (111.19-113.55) | *OAS1-2-3* |
|  | 6 | 2.36 | 1.45 | 33.99 (33.68-34.44) | *MHC* |
|  | 17 | 2.25 | 1.40 | 37.08 (36.93-37.76) | *HNF1B* |
|  | 6 | 2.17 | 0.55 | 25.00 (24.95-25.33) | *MHC* |
| AFR | 16 | 6.71 | 2.65 | 17.24 (16.75-18.01) | *XYLT1* |
|  | 19 | 4.44 | 2.45 | 17.25 (15.51-20.98) | . |
|  | 7 | 4.07 | 2.00 | 80.70 (80.13-81.34) | *SEMA3C* |
|  | 11 | 4.06 | 2.05 | 5.75 (5.27-6.32) | *HBB* |
|  | 9 | 3.45 | 1.25 | 134.60 (134.55-134.66) | . |
|  | 1 | 3.38 | 1.45 | 240.54 (240.45-240.64) | . |
|  | 19 | 3.21 | 1.20 | 3.09 (3.02-3.14) | . |
|  | 17 | 3.20 | 1.10 | 3.87 (3.80-3.93) | . |
|  | 11 | 3.09 | 0.90 | 19.94 (19.89-20.00) | . |
|  | 3 | 3.03 | 1.10 | 3.09 (3.04-3.09) | . |

**Table S6.**
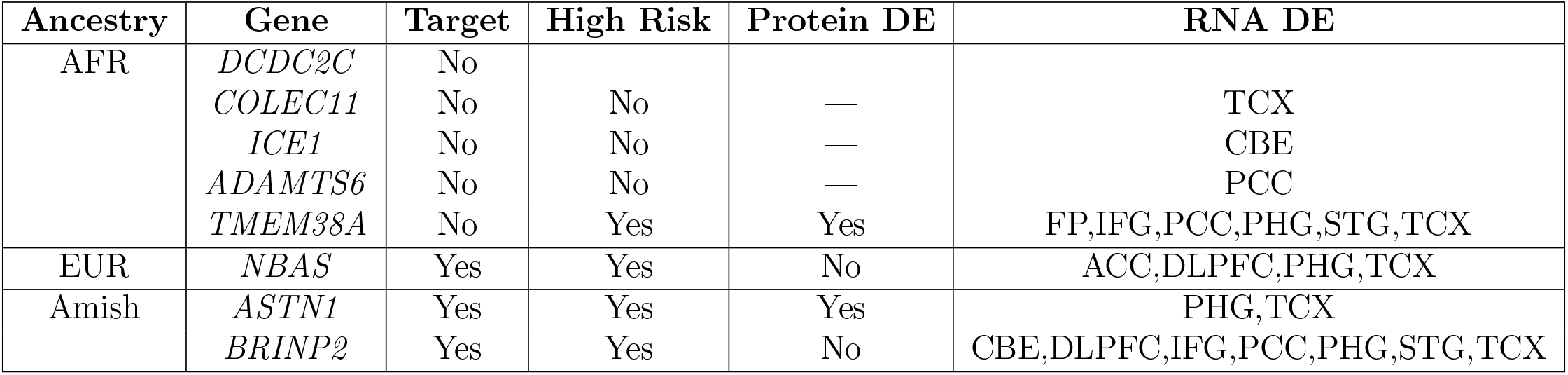
Annotations from Agora web resource of genes in risk loci. Each gene is located within or less than a few hundred kilobases of genome-wide significant loci in the case-control scans of three ancestry cohorts. Columns concern if the gene is a nominated therapeutic target (Target), has a multi-omic risk score greater than 3.80 (High Risk), has evidence of differential protein expression in post-mortem AD individuals, or has RNA differential expression (DE) in a brain region. The brain regions are the anterior cingulate cortex (ACC), cerebellum (CBE), dorsolateral prefrontal cortex (DLPFC), frontal pole (FP), inferior frontal gyrus (IFG), posterior cingulate cortex (PCC), parahippocampal gyrus (PHG), superior temporal gyrus (STG), and temporal cortex (TCX). Horizontal lines indicate that there is no data available for that measurement.

| Ancestry | Gene | Target | High Risk | Protein DE | RNA DE |
| --- | --- | --- | --- | --- | --- |
| AFR | *DCDC2C* | No | — | — | — |
|  | *COLEC11* | No | No | — | TCX |
|  | *ICE1* | No | No | — | CBE |
|  | *ADAMTS6* | No | No | — | PCC |
|  | *TMEM38A* | No | Yes | Yes | FP,IFG,PCC,PHG,STG,TCX |
| EUR | *NBAS* | Yes | Yes | No | ACC,DLPFC,PHG,TCX |
| Amish | *ASTN1* | Yes | Yes | Yes | PHG,TCX |
|  | *BRINP2* | Yes | Yes | No | CBE,DLPFC,IFG,PCC,PHG,STG,TCX |

