## Supplemental Acknowledgments for "Multiple-testing corrections in case-control studies using identity-by-descent segments"

### Supplementary Acknowledgements

Data for this study were prepared, archived, and distributed by the National Institute on Aging Alzheimer's Disease Data Storage Site (NIAGADS) at the University of Pennsylvania (U24-AG041689), funded by the National Institute on Aging.

Below, we provide acknowledgments for the specific datasets in the ADSP release 4 whole genome sequence (WGS) data. We accessed these study-specific acknowledgments at https://dss.niagads.org/datasets/ng00067/ on June 3, 2025.

**Alzheimer's Disease Sequencing Project (sa000001) data:**

The Alzheimer's Disease Sequencing Project (ADSP) comprises two Alzheimer's Disease (AD) genetics consortia and three National Human Genome Research Institute (NHGRI)- funded Large-Scale Sequencing and Analysis Centers (LSAC). The two AD genetics consortia are the Alzheimer's Disease Genetics Consortium (ADGC) funded by NIA (U01 AG032984), and the Cohorts for Heart and Aging Research in Genomic Epidemiology (CHARGE) funded by NIA (R01 AG033193), the National Heart, Lung, and Blood Institute (NHLBI), other National Institute of Health (NIH) institutes and other foreign governmental and non-governmental organizations. The Discovery Phase analysis of sequence data is supported through UF1AG047133 (to Drs. Schellenberg, Farrer, Pericak-Vance, Mayeux, and Haines); U01AG049505 to Dr. Seshadri; U01AG049506 to Dr. Boerwinkle; U01AG049507 to Dr. Wijsman; and U01AG049508 to Dr. Goate and the Discovery Extension Phase analysis is supported through U01AG052411 to Dr. Goate, U01AG052410 to Dr. Pericak-Vance and U01 AG052409 to Drs. Seshadri and Fornage.

Sequencing for the Follow Up Study (FUS) is supported through U01AG057659 (to Drs. Pericak-Vance, Mayeux, and Vardarajan) and U01AG062943 (to Drs. Pericak-Vance and Mayeux). Data generation and harmonization in the Follow-up Phase is supported by U54AG052427 (to Drs. Schellenberg and Wang). The FUS Phase analysis of sequence data is supported through U01AG058589 (to Drs. Destefano, Boerwinkle, De Jager, Fornage, Seshadri, and Wijsman), U01AG058654 (to Drs. Haines, Bush, Farrer, Martin, and Pericak-Vance), U01AG058635 (to Dr. Goate), RF1AG058066 (to Drs. Haines, Pericak-Vance, and Scott), RF1AG057519 (to Drs. Farrer and Jun), R01AG048927 (to Dr. Farrer), and RF1AG054074 (to Drs. Pericak-Vance and Beecham).

The ADGC cohorts include: Adult Changes in Thought (ACT) (U01 AG006781, U19 AG066567), the Alzheimer's Disease Research Centers (ADRC) (P30 AG062429, P30 AG066468, P30 AG062421, P30 AG066509, P30 AG066514, P30 AG066530, P30 AG066507, P30 AG066444, P30 AG066518, P30 AG066512, P30 AG066462, P30 AG072979, P30 AG072972, P30 AG072976, P30 AG072975, P30 AG072978, P30 AG072977, P30 AG066519, P30 AG062677, P30 AG079280, P30 AG062422, P30 AG066511, P30 AG072946, P30 AG062715, P30 AG072973, P30 AG066506, P30 AG066508, P30 AG066515, P30 AG072947,

P30 AG072931, P30 AG066546, P20 AG068024, P20 AG068053, P20 AG068077, P20 AG068082, P30 AG072958, P30 AG072959), the Chicago Health and Aging Project (CHAP) (R01 AG11101, RC4 AG039085, K23 AG030944), Indiana Memory and Aging Study (IMAS) (R01 AG019771), Indianapolis Ibadan (R01 AG009956, P30 AG010133), the Memory and Aging Project (MAP) ( R01 AG17917), Mayo Clinic (MAYO) (R01 AG032990, U01 AG046139, R01 NS080820, RF1 AG051504, P50 AG016574), Mayo Parkinson's Disease controls (NS039764, NS071674, 5RC2HG005605), University of Miami (R01 AG027944, R01 AG028786, R01 AG019085, IIRG09133827, A2011048), the Multi-Institutional Research in Alzheimer's Genetic Epidemiology Study (MIRAGE) (R01 AG09029, R01 AG025259), the National Centralized Repository for Alzheimer's Disease and Related Dementias (NCRAD) (U24 AG021886), the National Institute on Aging Late Onset Alzheimer's Disease Family Study (NIA- LOAD) (U24 AG056270), the Religious Orders Study (ROS) (P30 AG10161, R01 AG15819), the Texas Alzheimer's Research and Care Consortium (TARCC) (funded by the Darrell K Royal Texas Alzheimer's Initiative), Vanderbilt University/Case Western Reserve University (VAN/CWRU) (R01 AG019757, R01 AG021547, R01 AG027944, R01 AG028786, P01 NS026630, and Alzheimer's Association), the Washington Heights-Inwood Columbia Aging Project (WHICAP) (RF1 AG054023), the University of Washington Families (VA Research Merit Grant, NIA: P50AG005136, R01AG041797, NINDS: R01NS069719), the Columbia University Hispanic Estudio Familiar de Influencia Genetica de Alzheimer (EFIGA) (RF1 AG015473), the University of Toronto (UT) (funded by Wellcome Trust, Medical Research Council, Canadian Institutes of Health Research), and Genetic Differences (GD) (R01 AG007584). The CHARGE cohorts are supported in part by National Heart, Lung, and Blood Institute (NHLBI) infrastructure grant HL105756 (Psaty), RC2HL102419 (Boerwinkle) and the neurology working group is supported by the National Institute on Aging (NIA) R01 grant AG033193.

The CHARGE cohorts participating in the ADSP include the following: the Austrian Stroke Prevention Study (ASPS), ASPS-Family study, and the Prospective Dementia Registry-Austria (ASPS/PRODEM-Aus), the Atherosclerosis Risk in Communities (ARIC) Study, the Cardiovascular Health Study (CHS), the Erasmus Rucphen Family Study (ERF), the Framingham Heart Study (FHS), and the Rotterdam Study (RS). ASPS is funded by the Austrian Science Fund (FWF) grants P20545-P05 and P13180, as well as the Medical University of Graz. The ASPS-Fam is funded by the Austrian Science Fund (FWF) project I904, the EU Joint Programme – Neurodegenerative Disease Research (JPND) in the frame of the BRIDGET project (Austria, Ministry of Science), the Medical University of Graz, and the Steiermärkische Krankenanstalten Gesellschaft. PRODEM-Austria is supported by the Austrian Research Promotion agency (FFG) (Project No. 827462) and by the Austrian National Bank (Anniversary Fund, project 15435. ARIC research is carried out as a collaborative study supported by NHLBI contracts (HHSN268201100005C, HHSN268201100006C, HHSN268201100007C, HHSN268201100008C, HHSN268201100009C, HHSN268201100010C, HHSN268201100011C, and HHSN268201100012C).

Neurocognitive data in ARIC is collected by U01 2U01HL096812, 2U01HL096814, 2U01HL096899, 2U01HL096902, 2U01HL096917 from the NIH (NHLBI, NINDS, NIA and NIDCD), and with previous brain MRI examinations funded by R01-HL70825 from the NHLBI. CHS research was supported by contracts HHSN268201200036C, HHSN268200800007C, N01HC55222, N01HC85079, N01HC85080, N01HC85081, N01HC85082, N01HC85083, N01HC85086, and grants U01HL080295 and U01HL130114 from the NHLBI with additional contribution from the National Institute of Neurological Disorders and Stroke (NINDS). Additional support was provided by R01AG023629, R01AG15928, and R01AG20098 from the NIA. FHS research is supported by NHLBI contracts N01-HC-25195 and HHSN268201500001I. This study was also supported by additional grants from the NIA (R01s AG054076, AG049607 and AG033040 and NINDS (R01 NS017950). The ERF study as a part of EUROSPAN (European Special Populations Research Network) was supported by European Commission FP6 STRP grant number 018947 (LSHG-CT-2006- 01947) and also received funding from the European Community's Seventh Framework Programme (FP7/2007-2013)/grant agreement HEALTH-F4- 2007-201413 by the European Commission under the programme "Quality of Life and Management of the Living Resources" of 5th Framework Programme (no. QLG2-CT-2002-01254). High-throughput analysis of the ERF data was supported by a joint grant from the Netherlands Organization for Scientific Research and the Russian Foundation for Basic Research (NWO-RFBR 047.017.043). The Rotterdam Study is funded by Erasmus Medical Center and Erasmus University, Rotterdam, the Netherlands Organization for Health Research and Development (ZonMw), the Research Institute for Diseases in the Elderly (RIDE), the Ministry of Education, Culture and Science, the Ministry for Health, Welfare and Sports, the European Commission (DG XII), and the municipality of Rotterdam. Genetic data sets are also supported by the Netherlands Organization of Scientific Research NWO Investments (175.010.2005.011, 911-03-012), the Genetic Laboratory of the Department of Internal Medicine, Erasmus MC, the Research Institute for Diseases in the Elderly (014- 93-015; RIDE2), and the Netherlands Genomics Initiative (NGI)/Netherlands Organization for Scientific Research (NWO) Netherlands Consortium for Healthy Aging (NCHA), project 050-060-810. All the studies are grateful to their participants, faculty, and staff. The content of these manuscripts is solely the responsibility of the authors. It does not necessarily represent the official views of the National Institutes of Health or the U.S. Department of Health and Human Services.

The FUS cohorts include: the Alzheimer's Disease Research Centers (ADRC) (P30 AG062429, P30 AG066468, P30 AG062421, P30 AG066509, P30 AG066514, P30 AG066530, P30 AG066507, P30 AG066444, P30 AG066518, P30 AG066512, P30 AG066462, P30 AG072979, P30 AG072972, P30 AG072976, P30 AG072975, P30 AG072978, P30 AG072977, P30 AG066519, P30 AG062677, P30 AG079280, P30 AG062422, P30 AG066511, P30 AG072946, P30 AG062715, P30 AG072973, P30 AG066506, P30 AG066508, P30 AG066515, P30 AG072947, P30 AG072931, P30 AG066546, P20 AG068024, P20 AG068053, P20 AG068077, P20 AG068082, P30 AG072958, P30 AG072959), Alzheimer's Disease Neuroimaging Initiative (ADNI) (U19AG024904), Amish Protective Variant Study (RF1AG058066), Cache County

Study (R01AG11380, R01AG031272, R01AG21136, RF1AG054052), Case Western Reserve University Brain Bank (CWRUBB) (P50AG008012), Case Western Reserve University Rapid Decline (CWRURD) (RF1AG058267, NU38CK000480), CubanAmerican Alzheimer's Disease Initiative (CuAADI) (3U01AG052410), Estudio Familiar de Influencia Genetica en Alzheimer (EFIGA) (5R37AG015473, RF1AG015473, R56AG051876), Genetic and Environmental Risk Factors for Alzheimer Disease Among African Americans Study (GenerAAtions) (2R01AG09029, R01AG025259, 2R01AG048927), Gwangju Alzheimer and Related Dementias Study (GARD) (U01AG062602), Hillblom Aging Network (2014-A-004- NET, R01AG032289, R01AG048234), Hussman Institute for Human Genomics Brain Bank (HIHGBB) (R01AG027944, Alzheimer's Association "Identification of Rare Variants in Alzheimer Disease"), Ibadan Study of Aging (IBADAN) (5R01AG009956), Longevity Genes Project (LGP) and LonGenity (R01AG042188, R01AG044829, R01AG046949, R01AG057909, R01AG061155, P30AG038072), Mexican Health and Aging Study (MHAS) (R01AG018016), Multi-Institutional Research in Alzheimer's Genetic Epidemiology (MIRAGE) (2R01AG09029, R01AG025259, 2R01AG048927), Northern Manhattan Study (NOMAS) (R01NS29993), Peru Alzheimer's Disease Initiative (PeADI) (RF1AG054074), Puerto Rican 1066 (PR1066) (Wellcome Trust (GR066133/GR080002), European Research Council (340755)), Puerto Rican Alzheimer Disease Initiative (PRADI) (RF1AG054074), Reasons for Geographic and Racial Differences in Stroke (REGARDS) (U01NS041588), Research in African American Alzheimer Disease Initiative (REAAADI) (U01AG052410), the Religious Orders Study (ROS) (P30 AG10161, P30 AG72975, R01 AG15819, R01 AG42210), the RUSH Memory and Aging Project (MAP) (R01 AG017917, R01 AG42210Stanford Extreme Phenotypes in AD (R01AG060747), University of Miami Brain Endowment Bank (MBB), University of Miami/Case Western/North Carolina A&T African American (UM/CASE/NCAT) (U01AG052410, R01AG028786), and Wisconsin Registry for Alzheimer's Prevention (WRAP) (R01AG027161 and R01AG054047).

The four LSACs are: the Human Genome Sequencing Center at the Baylor College of Medicine (U54 HG003273), the Broad Institute Genome Center (U54HG003067), the American Genome Center at the Uniformed Services University of the Health Sciences (U01AG057659), and the Washington University Genome Institute (U54HG003079). Genotyping and sequencing for the ADSP FUS were also conducted at the John P. Hussman Institute for Human Genomics (HIHG) Center for Genome Technology (CGT).

Biological samples and associated phenotypic data used in primary data analyses were stored at the Study Investigators' institutions, and at the National Centralized Repository for Alzheimer's Disease and Related Dementias (NCRAD, U24AG021886) at Indiana University, funded by NIA. Associated Phenotypic Data used in primary and secondary data analyses were provided by Study Investigators, the NIA funded Alzheimer's Disease Centers (ADCs), and the National Alzheimer's Coordinating Center (NACC, U24AG072122) and the National Institute on Aging Genetics of Alzheimer's Disease Data Storage Site (NIAGADS, U24AG041689) at the University of Pennsylvania, funded by NIA. Harmonized phenotypes were provided by the

ADSP Phenotype Harmonization Consortium (ADSP-PHC), which is funded by the NIA (U24 AG074855, U01 AG068057, and R01 AG059716) and Ultrascale Machine Learning to Empower Discovery in Alzheimer's Disease Biobanks (AI4AD, U01 AG068057). This research was supported in part by the Intramural Research Program of the National Institutes of Health, National Library of Medicine.

Contributors to the Genetic Analysis Data included Study Investigators on projects that were individually funded by the NIA, as well as other NIH institutes, and by private U.S. organizations, or foreign governmental or nongovernmental organizations.

The ADSP Phenotype Harmonization Consortium (ADSP-PHC) is funded by NIA (U24 AG074855, U01 AG068057, and R01 AG059716). The harmonized cohorts within the ADSP-PHC include: the Anti-Amyloid Treatment in Asymptomatic Alzheimer's study (A4 Study), a secondary prevention trial in preclinical Alzheimer's disease, aiming to slow cognitive decline associated with brain amyloid accumulation in clinically normal older participants. The A4 Study is funded by a public-private-philanthropic partnership, including funding from the National Institutes of Health's National Institute on Aging, Eli Lilly and Company, the Alzheimer's Association, the Accelerating Medicines Partnership, the GHR Foundation, an anonymous foundation, and additional private donors, with in-kind support from Avid and Cogstate. The companion observational Longitudinal Evaluation of Amyloid Risk and Neurodegeneration (LEARN) Study is funded by the Alzheimer's Association and GHR Foundation. The A4 and LEARN Studies are led by Dr. Reisa Sperling at Brigham and Women's Hospital, Harvard Medical School and Dr. Paul Aisen at the Alzheimer's Therapeutic Research Institute (ATRI), University of Southern California. ATRI coordinates the A4 and LEARN Studies at the University of Southern California, and the data are made available through the Laboratory for Neuro Imaging at the University of Southern California. The participants in the A4 Study provided permission to share their de-identified data to advance the quest for a successful treatment for Alzheimer's disease. We acknowledge the dedication of all participants, site personnel, and partnership team members who continue to make the A4 and LEARN Studies possible. The complete A4 Study Team list is available on: a4study.org/a4-study-team.; the Adult Changes in Thought study (ACT), U01 AG006781, U19 AG066567; Alzheimer's Disease Neuroimaging Initiative (ADNI): Data collection and sharing for this project was funded by the Alzheimer's Disease Neuroimaging Initiative (ADNI) (National Institutes of Health Grant U01 AG024904) and DOD ADNI (Department of Defense award number W81XWH-12-2- 0012). ADNI is funded by the National Institute on Aging, the National Institute of Biomedical Imaging and Bioengineering, and through generous contributions from the following: AbbVie, Alzheimer's Association; Alzheimer's Drug Discovery Foundation; Araclon Biotech; BioClinica, Inc.; Biogen; Bristol- Myers Squibb Company; CereSpir, Inc.; Cogstate; Eisai Inc.; Elan Pharmaceuticals, Inc.; Eli Lilly and Company; EuroImmun; F. Hoffmann-La Roche Ltd and its affiliated company Genentech, Inc.; Fujirebio; GE Healthcare; IXICO Ltd.;Janssen Alzheimer Immunotherapy Research & Development, LLC.; Johnson & Johnson Pharmaceutical Research

& Development LLC.; Lumosity; Lundbeck; Merck & Co., Inc.;Meso Scale Diagnostics, LLC.; NeuroRx Research; Neurotrack Technologies; Novartis Pharmaceuticals Corporation; Pfizer Inc.; Piramal Imaging; Servier; Takeda Pharmaceutical Company; and Transition Therapeutics. The Canadian Institutes of Health Research is providing funds to support ADNI clinical sites in Canada. Private sector contributions are facilitated by the Foundation for the National Institutes of Health (www.fnih.org). The grantee organization is the Northern California Institute for Research and Education, and the study is coordinated by the Alzheimer's Therapeutic Research Institute at the University of Southern California. ADNI data are disseminated by the Laboratory for Neuro Imaging at the University of Southern California; Estudio Familiar de Influencia Genetica en Alzheimer (EFIGA): 5R37AG015473, RF1AG015473, R56AG051876; Memory & Aging Project at Knight Alzheimer's Disease Research Center (MAP at Knight ADRC): The Memory and Aging Project at the Knight-ADRC (Knight-ADRC). This work was supported by the National Institutes of Health (NIH) grants R01AG064614, R01AG044546, RF1AG053303, RF1AG058501, U01AG058922 and R01AG064877 to Carlos Cruchaga. The recruitment and clinical characterization of research participants at Washington University was supported by NIH grants P30AG066444, P01AG03991, and P01AG026276. Data collection and sharing for this project was supported by NIH grants RF1AG054080, P30AG066462, R01AG064614 and U01AG052410. We thank the contributors who collected samples used in this study, as well as patients and their families, whose help and participation made this work possible. This work was supported by access to equipment made possible by the Hope Center for Neurological Disorders, the Neurogenomics and Informatics Center (NGI: https://neurogenomics.wustl.edu/) and the Departments of Neurology and Psychiatry at Washington University School of Medicine; National Alzheimer's Coordinating Center (NACC): The NACC database is funded by NIA/NIH Grant U24 AG072122. NACC data are contributed by the NIA-funded ADRCs: P30 AG062429 (PI James Brewer, MD, PhD), P30 AG066468 (PI Oscar Lopez, MD), P30 AG062421 (PI Bradley Hyman, MD, PhD), P30 AG066509 (PI Thomas Grabowski, MD), P30 AG066514 (PI Mary Sano, PhD), P30 AG066530 (PI Helena Chui, MD), P30 AG066507 (PI Marilyn Albert, PhD), P30 AG066444 (PI John Morris, MD), P30 AG066518 (PI Jeffrey Kaye, MD), P30 AG066512 (PI Thomas Wisniewski, MD), P30 AG066462 (PI Scott Small, MD), P30 AG072979 (PI David Wolk, MD), P30 AG072972 (PI Charles DeCarli, MD), P30 AG072976 (PI Andrew Saykin, PsyD), P30 AG072975 (PI David Bennett, MD), P30 AG072978 (PI Neil Kowall, MD), P30 AG072977 (PI Robert Vassar, PhD), P30 AG066519 (PI Frank LaFerla, PhD), P30 AG062677 (PI Ronald Petersen, MD, PhD), P30 AG079280 (PI Eric Reiman, MD), P30 AG062422 (PI Gil Rabinovici, MD), P30 AG066511 (PI Allan Levey, MD, PhD), P30 AG072946 (PI Linda Van Eldik, PhD), P30 AG062715 (PI Sanjay Asthana, MD, FRCP), P30 AG072973 (PI Russell Swerdlow, MD), P30 AG066506 (PI Todd Golde, MD, PhD), P30 AG066508 (PI Stephen Strittmatter, MD, PhD), P30 AG066515 (PI Victor Henderson, MD, MS), P30 AG072947 (PI Suzanne Craft, PhD), P30 AG072931 (PI Henry Paulson, MD, PhD), P30 AG066546 (PI Sudha Seshadri, MD), P20 AG068024 (PI Erik Roberson, MD, PhD), P20 AG068053 (PI Justin Miller, PhD), P20 AG068077 (PI Gary Rosenberg, MD), P20 AG068082

(PI Angela Jefferson, PhD), P30 AG072958 (PI Heather Whitson, MD), P30 AG072959 (PI James Leverenz, MD); National Institute on Aging Alzheimer's Disease Family Based Study (NIA-AD FBS): U24 AG056270; Religious Orders Study (ROS): P30AG10161,R01AG15819, R01AG42210; Memory and Aging Project (MAP - Rush): R01AG017917, R01AG42210; Minority Aging Research Study (MARS): R01AG22018, R01AG42210; Washington Heights/Inwood Columbia Aging Project (WHICAP): RF1 AG054023;and Wisconsin Registry for Alzheimer's Prevention (WRAP): R01AG027161 and R01AG054047. Additional acknowledgments include the National Institute on Aging Genetics of Alzheimer's Disease Data Storage Site (NIAGADS, U24AG041689) at the University of Pennsylvania, funded by NIA.

**Alzheimer's Disease Neuroimaging Initiative (sa000002) data:**

Data collection and sharing for this project was funded by the Alzheimer's Disease Neuroimaging Initiative (ADNI) (National Institutes of Health Grant U01 AG024904) and DOD ADNI (Department of Defense award number W81XWH-12-2-0012). ADNI is funded by the National Institute on Aging, the National Institute of Biomedical Imaging and Bioengineering, and through generous contributions from the following: AbbVie, Alzheimer's Association; Alzheimer's Drug Discovery Foundation; Araclon Biotech; BioClinica, Inc.; Biogen; Bristol-Myers Squibb Company; CereSpir, Inc.; Cogstate; Eisai Inc.; Elan Pharmaceuticals, Inc.; Eli Lilly and Company; EuroImmun; F. Hoffmann-La Roche Ltd and its affiliated company Genentech, Inc.; Fujirebio; GE Healthcare; IXICO Ltd.; Janssen Alzheimer Immunotherapy Research & Development, LLC.; Johnson & Johnson Pharmaceutical Research & Development LLC.; Lumosity; Lundbeck; Merck & Co., Inc.; Meso Scale Diagnostics, LLC.; NeuroRx Research; Neurotrack Technologies; Novartis Pharmaceuticals Corporation; Pfizer Inc.; Piramal Imaging; Servier; Takeda Pharmaceutical Company; and Transition Therapeutics. The Canadian Institutes of Health Research is providing funds to support ADNI clinical sites in Canada. Private sector contributions are facilitated by the Foundation for the National Institutes of Health (www.fnih.org). The grantee organization is the Northern California Institute for Research and Education, and the study is coordinated by the Alzheimer's Therapeutic Research Institute at the University of Southern California. ADNI data are disseminated by the Laboratory for Neuro Imaging at the University of Southern California.

**Alzheimer's Disease Genetics Consortium (sa000003) data:**

The Alzheimer's Disease Genetics Consortium (ADGC) supported sample preparation, sequencing and data processing through NIA grant U01AG032984. Sequencing data generation and harmonization is supported by the Genome Center for Alzheimer's Disease, U54AG052427, and data sharing is supported by NIAGADS, U24AG041689. Samples from the National Centralized Repository for Alzheimer's Disease and Related Dementias (NCRAD), which receives government support under a cooperative agreement grant (U24 AG021886) awarded by the National Institute on Aging (NIA), were used in this study. We thank the contributors who collected the samples used in this study, as well as the patients and their families, whose help and participation made this work possible.

**ADGC-TARCC-WGS (snd10030) data:**

This study was made possible by the Texas Alzheimer's Research and Care Consortium (TARCC) funded by the state of Texas through the Texas Council on Alzheimer's Disease and Related Disorders and the Darrell K Royal Texas Alzheimer's Initiative.

**The Familial Alzheimer Sequencing Project (sa000004) and the Charles F. and Joanne Knight Alzheimer's Disease Research Center (sa000008) data:**

This work was supported by grants from the National Institutes of Health (R01AG044546, P01AG003991, RF1AG053303, R01AG058501, U01AG058922, RF1AG058501 and R01AG057777). The recruitment and clinical characterization of research participants at Washington University were supported by NIH P50 AG05681, P01 AG03991, and P01 AG026276. This work was supported by access to equipment made possible by the Hope Center for Neurological Disorders and the Departments of Neurology and Psychiatry at Washington University School of Medicine.

We thank the contributors who collected samples used in this study, as well as patients and their families, whose help and participation made this work possible. This work was supported by access to equipment made possible by the Hope Center for Neurological Disorders and the Departments of Neurology and Psychiatry at Washington University School of Medicine.

**ADSP-PHC harmonized phenotypes deposited within dataset, ng00067:**

The Memory and Aging Project at the Knight-ADRC (Knight-ADRC), supported by NIH grants R01AG064614, R01AG044546, RF1AG053303, RF1AG058501, U01AG058922 and R01AG064877 to Carlos Cruchaga. The recruitment and clinical characterization of research participants at Washington University was supported by NIH grants P30AG066444, P01AG03991, and P01AG026276. Data collection and sharing for this project were supported by NIH grants RF1AG054080, P30AG066462, R01AG064614, and U01AG052410. This work was supported by access to equipment made possible by the Hope Center for Neurological Disorders, the Neurogenomics and Informatics Center (NGI: https://neurogenomics.wustl.edu/), and the Departments of Neurology and Psychiatry at Washington University School of Medicine.

**Accelerating Medicines Partnership-Alzheimer's Disease (AMP-AD) (sa000011) data:**

Mayo RNAseq Study - Study data were provided by the following sources: The Mayo Clinic Alzheimer's Disease Genetic Studies, led by Dr. Nilufer Ertekin-Taner and Dr. Steven G. Younkin, Mayo Clinic, Jacksonville, FL using samples from the Mayo Clinic Study of Aging, the Mayo Clinic Alzheimer's Disease Research Center, and the Mayo Clinic Brain Bank. Data collection was supported through funding by NIA grants P50 AG016574, R01 AG032990, U01 AG046139, R01 AG018023, U01 AG006576, U01 AG006786, R01 AG025711, R01 AG017216,

R01 AG003949, NINDS grant R01 NS080820, CurePSP Foundation, and support from Mayo Foundation. Study data includes samples collected through the Sun Health Research Institute Brain and Body Donation Program of Sun City, Arizona. The Brain and Body Donation Program is supported by the National Institute of Neurological Disorders and Stroke (U24 NS072026 National Brain and Tissue Resource for Parkinson's Disease and Related Disorders), the National Institute on Aging (P30 AG19610 Arizona Alzheimer's Disease Core Center), the Arizona Department of Health Services (contract 211002, Arizona Alzheimer's Research Center), the Arizona Biomedical Research Commission (contracts 4001, 0011, 05-901 and 1001 to the Arizona Parkinson's Disease Consortium) and the Michael J. Fox Foundation for Parkinson's Research

ROSMAP - We are grateful to the participants in the Religious Order Study, the Memory and Aging Project (ROSMAP). This work was supported by the US National Institutes of Health [U01 AG046152, R01 AG043617, R01 AG042210, R01 AG036042, R01 AG036836, R01 AG032990, R01 AG18023, RC2 AG036547, P50 AG016574, U01 ES017155, KL2 RR024151, K25 AG041906-01, R01 AG30146, P30 AG10161, R01 AG17917, R01 AG15819, K08 AG034290, P30 AG10161 and R01 AG11101.

Mount Sinai Brain Bank (MSBB) - This work was supported by the grants R01AG046170, RF1AG054014, RF1AG057440 and R01AG057907 from the NIH/National Institute on Aging (NIA). R01AG046170 is a component of the AMP-AD Target Discovery and Preclinical Validation Project. Brain tissue collection and characterization was supported by NIH HHSN271201300031C.

**University of Pittsburg- Kamboh (sa000012) data:**

This study was supported by the National Institute on Aging (NIA) grants AG030653, AG041718, AG064877, and P30-AG066468.

**NACC Genentech Study (sa000013) data:**

We thank study participants, their families, and the sample collectors for their invaluable contributions. This research was supported in part by the National Institute on Aging grant U01AG049508 (PI Alison M. Goate). This research was supported in part by Genentech, Inc. (PI Alison M. Goate, Robert R. Graham).

The NACC database is funded by NIA/NIH Grant U01 AG016976. These NIA-funded ADCs contribute NACC data: P30 AG013846 (PI Neil Kowall, MD), P50 AG008702 (PI Scott Small, MD), P50 AG025688 (PI Allan Levey, MD, PhD), P30 AG010133 (PI Andrew Saykin, PsyD), P50 AG005146 (PI Marilyn Albert, PhD), P50 AG005134 (PI Bradley Hyman, MD, PhD), P50 AG016574 (PI Ronald Petersen, MD, PhD), P30 AG013854 (PI M. Marsel Mesulam, MD), P30 AG008017 (PI Jeffrey Kaye, MD), P30 AG010161 (PI David Bennett, MD), P30 AG010129 (PI

Charles DeCarli, MD), P50 AG016573 (PI Frank LaFerla, PhD), P50 AG005131 (PI Douglas Galasko, MD), P30 AG028383 (PI Linda Van Eldik, PhD), P30 AG010124 (PI John Trojanowski, MD, PhD), P50 AG005142 (PI Helena Chui, MD), P30 AG012300 (PI Roger Rosenberg, MD), P50 AG005136 (PI Thomas Grabowski, MD), P50 AG005681 (PI John Morris, MD), P30 AG028377 (Kathleen Welsh-Bohmer, PhD), and P50 AG008671 (PI Henry Paulson, MD, PhD).

Samples from the National Cell Repository for Alzheimer's Disease (NCRAD), which receives government support under a cooperative agreement grant (U24 AG21886) awarded by the National Institute on Aging (NIA), were used in this study. We thank the contributors who collected the samples used in this study, as well as the patients and their families, whose help and participation made this work possible.

The Alzheimer's Disease Genetics Consortium supported the collection of samples used in this study through National Institute on Aging (NIA) grants U01AG032984 and RC2AG036528.

**Cache County Study (sa000014) data:**

We acknowledge the generous contributions of the participants in the Cache County Memory Study. Sequencing for this study was funded by RF1AG054052 (PI: John S.K. Kauwe).

**NIH, CurePSP and Tau Consortium PSP WGS (sa000015) data:**

This project was funded by the NIH grant UG3NS104095 and supported by grants U54NS100693 and U54AG052427. Queen Square Brain Bank is supported by the Reta Lila Weston Institute for Neurological Studies and the Medical Research Council UK. The Mayo Clinic Florida had support from a Morris K. Udall Parkinson's Disease Research Center of Excellence (NINDS P50 #NS072187), CurePSP and the Tau Consortium. The samples from the University of Pennsylvania are supported by NIA grant P01AG017586.

**CurePSP and Tau Consortium PSP WGS (sa000016) data:**

This project was funded by the Tau Consortium, Rainwater Charitable Foundation, and CurePSP. It was also supported by NINDS grant U54NS100693 and NIA grants U54NS100693 and U54AG052427. Queen Square Brain Bank is supported by the Reta Lila Weston Institute for Neurological Studies and the Medical Research Council UK. The Mayo Clinic Florida had support from a Morris K. Udall Parkinson's Disease Research Center of Excellence (NINDS P50 #NS072187), CurePSP, and the Tau Consortium. The samples from the University of Pennsylvania are supported by NIA grant P01AG017586. Tissues were received from the Victorian Brain Bank, supported by The Florey Institute of Neuroscience and Mental Health, The Alfred, and the Victorian Forensic Institute of Medicine, and funded in part by Parkinson's Victoria and MND Victoria. We are grateful to the Sun Health Research Institute Brain and Body Donation Program in Sun City, Arizona, for providing human biological materials. The Brain and Body Donation Program is supported by the National Institute of Neurological Disorders and

Stroke (U24 NS072026 National Brain and Tissue Resource for Parkinson's Disease and Related Disorders), the National Institute on Aging (P30 AG19610 Arizona Alzheimer's Disease Core Center), the Arizona Department of Health Services ( contract 211002, Arizona Alzheimer's Research Center), the Arizona Biomedical Research Commission (contracts 4001, 0011, 05-901 and 1001 to the Arizona Parkinson's Disease Consortium) and the Michael J. Fox Foundation for Parkinson's Research. The biomaterial was provided by the Study Group DESCRIBE of the Clinical Research Unit at the German Center for Neurodegenerative Diseases (DZNE).

**UCLA Progressive Supranuclear Palsy (sa000017) data:**

We thank the AL-108-231 investigators for their contribution to this consortium dataset.

**The Diagnostic Assessment of Dementia for the Longitudinal Aging Study of India (LASI-DAD) (sa000019) data:**

The Longitudinal Aging Study in India, Diagnostic Assessment of Dementia data is sponsored by the National Institute on Aging (grant numbers R01AG051125 and U01AG065958) and is conducted by the University of Southern California.

**Dissecting the Genomic Etiology of non-Mendelian Early-Onset Alzheimer Disease (EOAD) and Related Phenotypes (sa000023) data:**

This work was supported by the National Institutes of Health (NIH) grant R01AG064614. The ADSP-FUS is supported by U01AG057659.

The National Institutes of Health, National Institute on Aging (NIH-NIA) supported this work through the following grants: ADGC, U01 AG032984, RC2 AG036528; samples from the National Centralized Repository for Alzheimer’s Disease and Related Dementias (NCRAD), which receives government support under a cooperative agreement grant (U24 AG21886) awarded by the National Institute on Aging (NIA), were used in this study. Sequencing data generation and harmonization are supported by the Genome Center for Alzheimer’s Disease, U54AG052427, and data sharing is supported by NIAGADS, U24AG041689. We thank the contributors who collected the samples used in this study, as well as the patients and their families, whose help and participation made this work possible.

NIH grants supported enrollment and data collection for the individual studies including the Alzheimer’s Disease Centers (ADC, P30 AG062429 (PI James Brewer, MD, PhD), P30 AG066468 (PI Oscar Lopez, MD), P30 AG062421 (PI Bradley Hyman, MD, PhD), P30 AG066509 (PI Thomas Grabowski, MD), P30 AG066514 (PI Mary Sano, PhD), P30 AG066530 (PI Helena Chui, MD), P30 AG066507 (PI Marilyn Albert, PhD), P30 AG066444 (PI John Morris, MD), P30 AG066518 (PI Jeffrey Kaye, MD), P30 AG066512 (PI Thomas Wisniewski, MD), P30 AG066462 (PI Scott Small, MD), P30 AG072979 (PI David Wolk, MD), P30 AG072972 (PI Charles DeCarli, MD), P30 AG072976 (PI Andrew Saykin, PsyD), P30 AG072975 (PI David Bennett, MD), P30 AG072978 (PI Neil Kowall, MD), P30 AG072977 (PI

Robert Vassar, PhD), P30 AG066519 (PI Frank LaFerla, PhD), P30 AG062677 (PI Ronald Petersen, MD, PhD), P30 AG079280 (PI Eric Reiman, MD), P30 AG062422 (PI Gil Rabinovici, MD), P30 AG066511 (PI Allan Levey, MD, PhD), P30 AG072946 (PI Linda Van Eldik, PhD), P30 AG062715 (PI Sanjay Asthana, MD, FRCP), P30 AG072973 (PI Russell Swerdlow, MD), P30 AG066506 (PI Todd Golde, MD, PhD), P30 AG066508 (PI Stephen Strittmatter, MD, PhD), P30 AG066515 (PI Victor Henderson, MD, MS), P30 AG072947 (PI Suzanne Craft, PhD), P30 AG072931 (PI Henry Paulson, MD, PhD), P30 AG066546 (PI Sudha Seshadri, MD), P20 AG068024 (PI Erik Roberson, MD, PhD), P20 AG068053 (PI Justin Miller, PhD), P20 AG068077 (PI Gary Rosenberg, MD), P20 AG068082 (PI Angela Jefferson, PhD), P30 AG072958 (PI Heather Whitson, MD), P30 AG072959 (PI James Leverenz, MD). The Miami ascertainment and research were supported in part through grants RF1AG054080, R01AG027944, R01AG019085, R01AG028786-02, and RC2AG036528. The Columbia ascertainment and research were supported in part through grants R37AG015473 and U24AG056270. The University of Washington's ascertainment and research were supported in part through grants R01AG044546, RF1AG053303, RF1AG058501, U01AG058922, and R01AG064877.
